# Immune-mediated indirect interaction between gut microbiota and bacterial pathogens

**DOI:** 10.1101/2025.03.31.646282

**Authors:** Maryam Keshavarz, Mathias Franz, Haicheng Xie, Caroline Zanchi, Susan Mbedi, Sarah Sparmann, Jens Rolff

**Author notes:** Corresponding author: Maryam Keshavarz.,. Contributing authors,.

## Abstract

**Background:** Organism survival depends on the ability to orchestrate interactions between the host immune system and gut microbiota in response to pathogenic infections. These tripartite interactions shape pathogen virulence evolution. A key regulator of the immune system and, hence, bipartite interactions (in insects) is the immune deficiency (Imd) pathway, which modulates gut microbiota and pathogens by synthesizing antimicrobial peptides (AMPs). However, whether Imd-dependent AMPs mediate indirect interactions between gut microbiota and pathogens in a tripartite context remains unclear. Using RNAi-mediated knockdown of *Tenebrio molitor Relish* (*TmRelish*), we hypothesized that Imd-dependent AMPs influence indirect interaction between *Providencia burhodogranaria_B* infection and the gut microbiota.

**Results:** While *TmRelish*-knockdown altered bipartite interactions, disrupting gut microbiota load and composition and increasing pathogen load and virulence through higher host mortality, it did not support our tripartite hypothesis. Instead of Imd-dependent AMPs, gut microbiota and pathogen were influenced by Imd-independent AMPs expression, suggesting alternative regulatory pathways. Nevertheless, our investigations of tripartite interactions showed a positive effect of *P. b_B* infection on gut microbiota load, while survival analysis further revealed a negative effect of gut microbiota on pathogens infection, suggesting microbiota- mediated immune priming.

**Conclusions:** These findings suggest that while Imd-dependent AMPs may not mediate tripartite interactions, microbiota-host interactions, such as microbiota-mediated immune priming and changes in microbiota load, can shape infection outcomes. These effects on infection outcomes almost certainly exert important selective pressures on the evolution of bacterial virulence.

## Background

Pathogen virulence and its evolutionary change can depend on the presence of host gut microbiota [1–3]. The gut microbiota, however, can be a double-edged sword, by either mediating protection [4–6], or facilitating infection by invading pathogens [7, 8] with opposite outcomes for host survival. Also, the metabolic [9, 10] and immunological environments [11, 12] are key contributors to the pathogenicity of invaders. These tripartite interactions between host, its microbiota and pathogens have so far gained little attention in invertebrates. One of the few studies showed in *Anopheles stephensi*, that fungal infection by *Beauveria bassiana* downregulated gut dual oxidase (Duox) and antimicrobial peptides (AMPs), leading to the overgrowth and translocation of the bacterium *Serratia marcescens* from the gut to the hemocoel, ultimately increasing host mortality [13]. Another example is found in the Pacific oyster, *Crassotrea gigas*, where viral infection of haemocytes by *Ostreid herpesvirus* impaired AMP expression, leading to a significant increase in the load of rare gut microbiota, such as *Vibrio* and *Arcobacter*, promoting host death [14]. Thus, in both cases, pathogen infections regulate AMP-mediated host defences, leading to alterations in gut microbiota load, suggesting that pathogens and gut microbiota interact indirectly through immune-mediated mechanisms.

Host immune-mediated indirect interactions between gut microbiota and pathogens could appear in different forms, based on the relative strength of effects established between gut microbiota and pathogens with their host immune system [1]. Depending on the balance between immune activation and suppression, host-pathogen-microbiota interactions can be negatively or positively modified and classified as apparent competition, indirect amensalism, or apparent mutualism [15]. In apparent competition, the negative effect could be symmetric for example when host immunity equally suppresses both microbiota and pathogens, or asymmetric when one is more strongly suppressed than the other. Indirect amensalism occurs when host immunity exerts a disproportionately negative effect on pathogen infection while allowing gut microbiota to thrive, thereby favouring the microbiota over the pathogen or vice versa. In contrast, apparent mutualism arises when host immunity has a positive effect, facilitating the proliferation of both microbiota and pathogens. Ultimately, the nature of these immune-mediated interactions imposes selection pressures on pathogen traits, potentially shaping virulence evolution [1, 15, 16]. Predictions for different forms of interactions in virulence evolution depend on such tripartite interactions [1, 15]. Earlier studies in mosquitoes and *Drosophila* demonstrated examples of apparent competition [17–19], indirect amensalism [20], or apparent mutualism [13, 14] between gut microbiota and pathogenic infections, mediated through host immunity.

Host immunity comprises multiple lines of defence that mediate tolerance toward gut microbiota (host-microbiota) while providing resistance against pathogen infections through antimicrobial activity (host-pathogen). One of the well-known innate signalling pathways in insects is the Imd pathway, which is considered a master regulator of gut microbiota and a key defence mechanism against pathogenic infections [21, 22]. Studies in *Drosophila*, for example, have demonstrated that local gut Imd-induced immunity confronts ingested microorganisms through synthesizing a cocktail of effector molecules, including AMPs, thereby regulating microbial community dynamics and host interactions [23, 24]. One of the key components of Imd pathway is the NF-κB transcription factor, Relish, that translocate into the nucleus to activate AMP expression [25]. This is supported by studies in *Relish*-deficient insects, i.e., disruption in AMPs expression, such as *Drosophila* [20, 21, 26], the red palm weevil (*Rhynchophorus ferrugineus*) [27], and the oriental fruit fly *Bactrocera dorsalis* [28], all of which exhibit changes in both the composition and bacterial loads of their gut-associated microbes. Yet, alterations in gut microbiota are not solely driven by changes in AMP expression. Gut microbiota-induced reactive oxygen species (ROS) regulate Imd-mediated AMP expression, suggesting a crosstalk between Duox and the pathway [26, 29]. In addition, there is compelling evidence in fruit flies [22] and mealworm beetles, *Tenebrio molitor* [30], that *Relish* knockdown renders insects more susceptible to pathogenic infections. Five families of inducible antibacterial AMPs with multiple isoforms have been identified in *T. molitor*: *Attacins* [31], *Tenecins* [32], *Coleoptericins* [33], *Defensins* [34], and *Cecropins* [35]. Here, we investigate the Imd-dependent AMP expression interaction between microbiota and pathogens in an insect host.

Given the importance of Relish-mediated AMPs in bipartite interactions between host-microbiota and host-pathogen relationships, we hypothesize that this pathway also plays a role in shaping indirect interactions (positive or negative effect) between pathogens and gut microbiota. In the present work, we used RNAi-mediated knockdown of *Relish* to generate different transcriptional levels of AMPs in the mealworm beetle, *Tenebrio molitor* [30]. As a bacterial model, we used *Providencia burhodogranariea_B* (*P. b_B*), a Gram-negative bacterium isolated from the hemolymph of wild *Drosophila melanogaster* [36].

To test our hypothesis, we first established how the bipartite interactions between the host and either its microbiota or a pathogen is influenced by Imd-mediated AMPs expression in our model system. We then investigated whether and which form of immune-mediated indirect interactions occur in tripartite interactions context. We addressed if Imd-mediated AMPs expression mediates (1) the impact of pathogen infection on gut microbiota (Fig. 1a) and predicted that the effect of pathogen infection on gut microbiota should be attenuated or absent in *Relish*-knockdown larvae (P1). Secondly, we studied the influence of gut microbiota on pathogen infection (Fig. 1b) and predicted also that the influence of gut microbiota on pathogen infection should be absent or reduced in *Relish*-knockdown larvae (P2).

**Fig 1.**
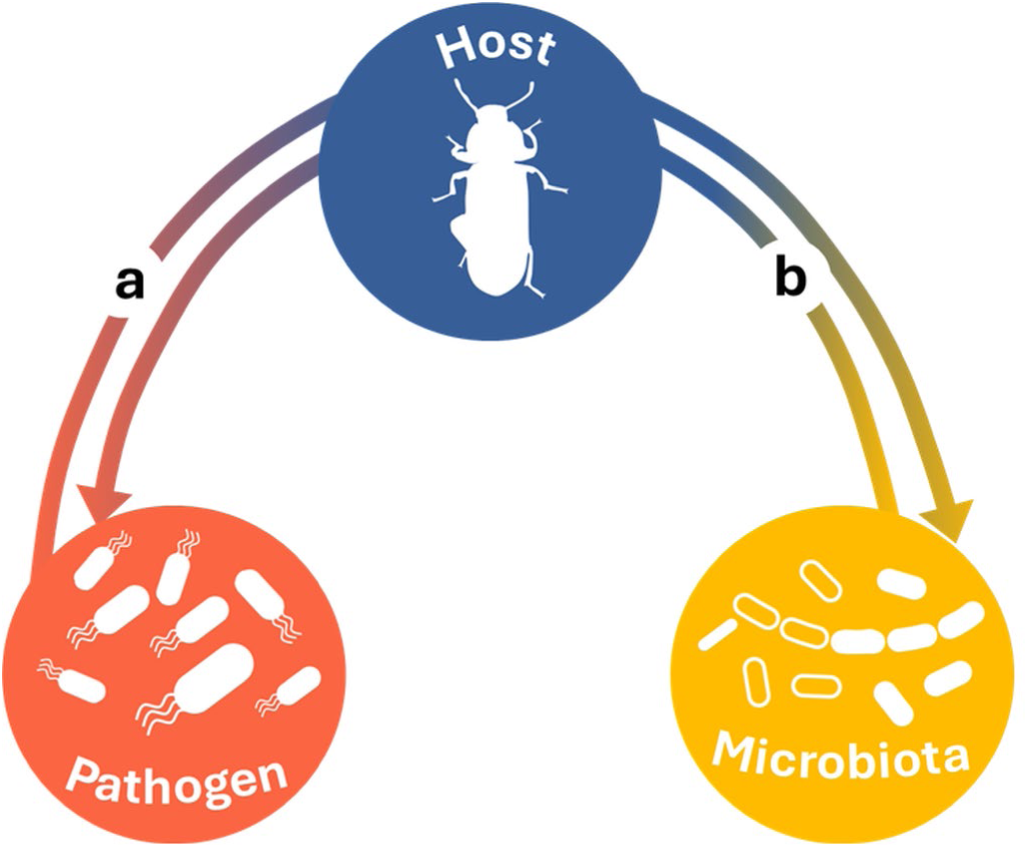
Indirect interaction between gut microbiota and pathogen via the host immune system. **(a)** Pathogenic infection affects the gut microbiota prevalence due to its effect on the host immune response (**b**) The gut microbiota affects pathogenic infection outcomes due to its effect on the host immune response.

## Results

### Bipartite interaction between host and gut microbiota

We found that the knockdown of *TmRelish* led to a significant increase in microbial load, as demonstrated by the higher CFUs in ds*TmRelish*-treated larvae compared to the ds*EGFP* group (z = 3.461, *p* < 0.001) (Fig. 2a). A Bray-Curtis dissimilarity matrix analysis based on differential bacterial genus presence and abundance revealed a significant effect of the knockdown treatment on bacterial microbiota composition (R² = 0.29, F = 8.52, *p* < 0.001), (Fig. 2b). Notably, the normalized counts of three operational taxonomic units (OTUs), including two assigned to the genus *Bacillus* (family *Bacillaceae 1* and *Bacillaceae 2*) and one to the genus *Pediococcus* (family *Lactobacillaceae*), were significantly reduced following *TmRelish* knockdown (adjusted *p* < 0.05) (Fig. 2 c–e). These findings indicate a regulatory role of *TmRelish* in maintaining a balanced gut microbial community, potentially through modulating the proliferation of specific gut bacterial taxa. 16S rRNA amplicon sequence analysis revealed that *Clostridiales*, *Entomoplasmatales*, *Lactobacillales*, *Bacillales*, and *Enterobacteriales* dominated the ds*EGFP*-treated larvae (Fig. 2f). In ds*TmRelish*-treated larvae, certain bacterial orders, such as *Entomoplasmatales* (genus: *Spiroplasma*), *Enterobacteriales*, and *Lactobacillales*, were dominant, while some were absent compared to ds*EGFP*-treated controls (Fig. S1); although *Spiroplasma* showed a visually notable increase (Fig. 2f), differential abundance analysis revealed this increase was not statistically significant (adjusted *p* < 0.1).

**Fig 2.**
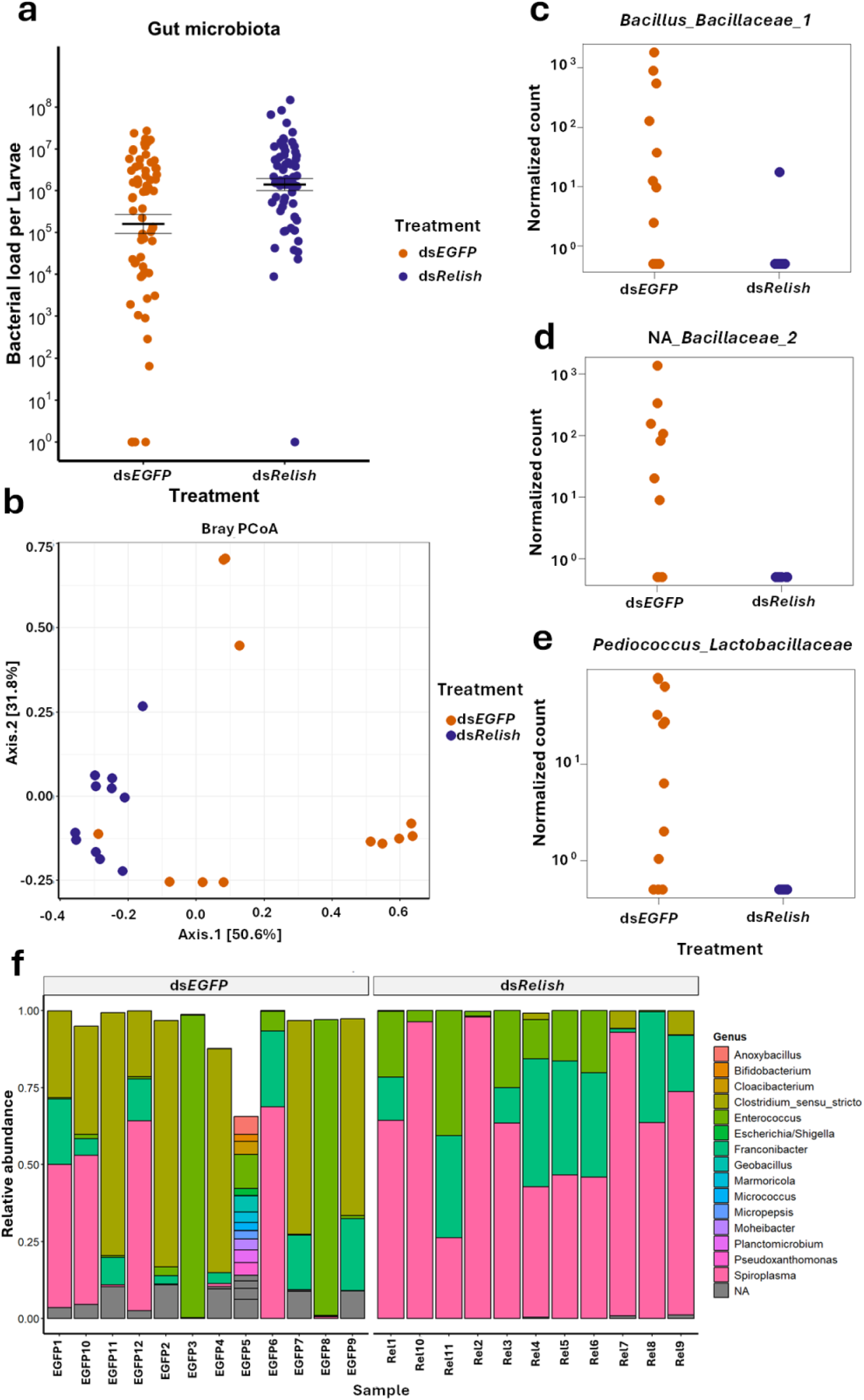
Bacterial load, microbiota composition, and differential abundance in larvae treated with ds*EGFP* and ds*TmRelish*. (**a**) Bacterial load of all culturable bacteria in *Tenebrio molitor* larvae treated with ds*EGFP* and ds*TmRelish* was quantified using trypticase soy agar (TSA) plates. Means and standard errors are represented by the black lines, while the dots represent the biological replicates from individual guts (n=∼12 per treatment, six biological replicates). (**b**) Principal coordinates analysis (PcoA) plot visualizing Bray-Curtis dissimilarity of the composition of the bacterial gut microbiota between ds*EGFP* (orange) and ds*TmRelish* (purple) *T. molitor* larvae. Each point represents the gut microbiota composition of an individual larva, with colour indicating the treatment group. The axis labels indicate the percentage of variation captured by each dimension. A PERMANOVA with 999 permutations showed that the treatment significantly separates the samples. (**c**) ASV counts of an OTU member of the *Bacillus* genus (family *Bacillaceae*_1), (**d**) ASV counts of an OTU belonging to an unidentified genus (family *Bacillaceae*_2), and (**e**) ASV counts of an OTU member of the *Pediococcus* genus (family *Lactobacillaceae*) illustrated between ds*EGFP* (orange) and ds*TmRelish* (purple) larvae. Each point represents a single larva, with colours indicating the treatment. (**f**) Relative abundance of the top 20 genera detected by 16S rRNA gene sequencing in *T. molitor* larvae treated with dsRNA, visualized by bar plots. Each bar represents an individual sample, with coloured box indicating different taxa. The hight of each box represents to the relative abundance of that taxon within the samples. Grey boxes indicate OTUs for which no taxonomy could be assigned.

### Bipartite interaction between host and pathogen

Next, we investigated whether *TmRelish* knockdown affects host survival and expression levels of AMPs following infection. When we scored survival, no death was observed among larvae treated with PBS, as well as in the control-*P. b_B* groups (CR-PBS, ds*EGFP*-PBS, ds*TmRelish*-PBS, and CR-*P. b_B*). Therefore, these groups were excluded from further analysis, however, the corresponding data are presented in the figure (Fig. 3a).

**Fig 3.**
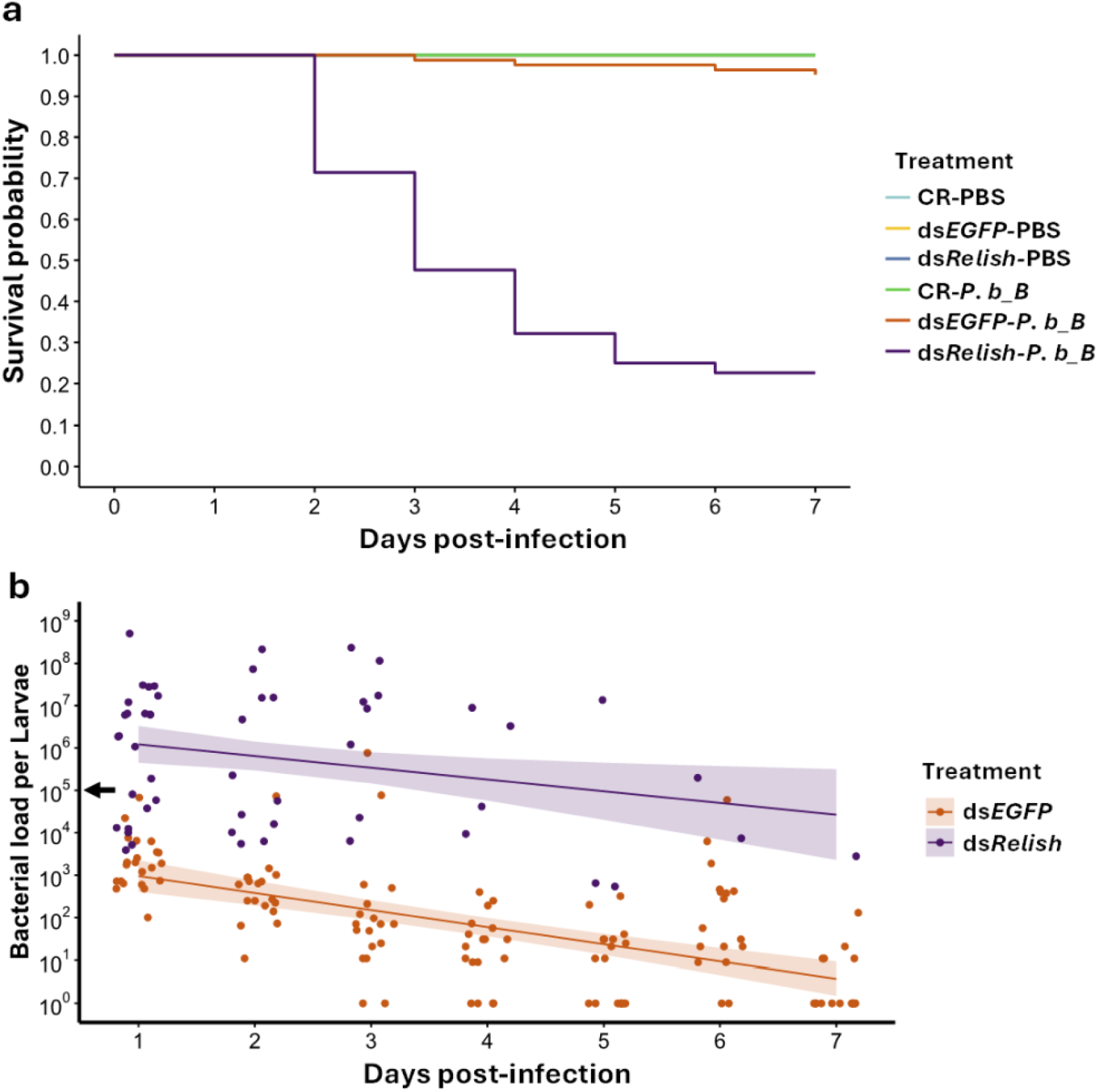
Host survival and *Providencia burhodogranariea_B* load in *TmRelish* knockdown larvae. (**a**) Survival of *T. molitor* larvae subjected to control (no dsRNA injection), ds*EGFP*, or ds*TmRelish* treatments, following exposure to either *P. burhodogranariea_B* (*P. b_B*) or PBS (n=∼30 per treatment, three independent experiments) was monitored over a period of seven days. (**b**) Bacterial load of *P. burhodogranariea_B* in ds*EGFP* and ds*TmRelish*-treated larvae over a period of seven days (n=∼10 per treatment per day, two independent experiments). The black arrow on the y-axis indicates the approximate injection dose. Purple and orange lines and associated bands depict the estimates and corresponding 95% confidence intervals of a linear model. *TmRelish* knockdown larvae were short-lived compared to *EGFP* controls (X² _1, 18_ = 50.80, *p* < 0.001), accompanied by an increased bacterial load (ds*EGFP*/ds*TmRelish*: F _1, 176_ = 232.09, *p* < 0.001).

We found that more ds*TmRelish* larvae had died on day 7 compared to ds*EGFP* larvae (X² _1, 18_ = 50.80, *p* < 0.001) (Fig. 3a). The bacterial load in both ds*EGFP*- and ds*TmRelish*-treated larvae decreased over the 7-day period (time: *F* _1, 176_ = 59.30, *p* < 0.001) (Fig. 3b). As expected, the *P. burhodogranariea_B* survival was enhanced in ds*TmRelish* compared to ds*EGFP* larvae over the 7-day period (ds*EGFP*/ds*TmRelish*: F _1, 176_ = 232.09, *p* < 0.001) (Fig. 3b).

When investigating AMP expression, we found a significant interaction between knockdown treatment and infection on *TmAtt1a*, *TmAtt1b*, *TmTen1, TmTen2 TmTen4*, *TmColA*, and *TmColB* expression levels (Fig. 4, Table S1). Infections generally led to an upregulation of these AMPs compared to non-infected PBS controls. However, for these seven AMPs this upregulation was stronger in ds*EGFP*-treated controls compared to ds*TmRelish*-treated larvae (Fig. 4). This suggests that *TmRelish* is essential for infection-induced *AMP* upregulation, while its knockdown has little effect on basal AMP expression in non-infected larvae.

**Fig 4.**
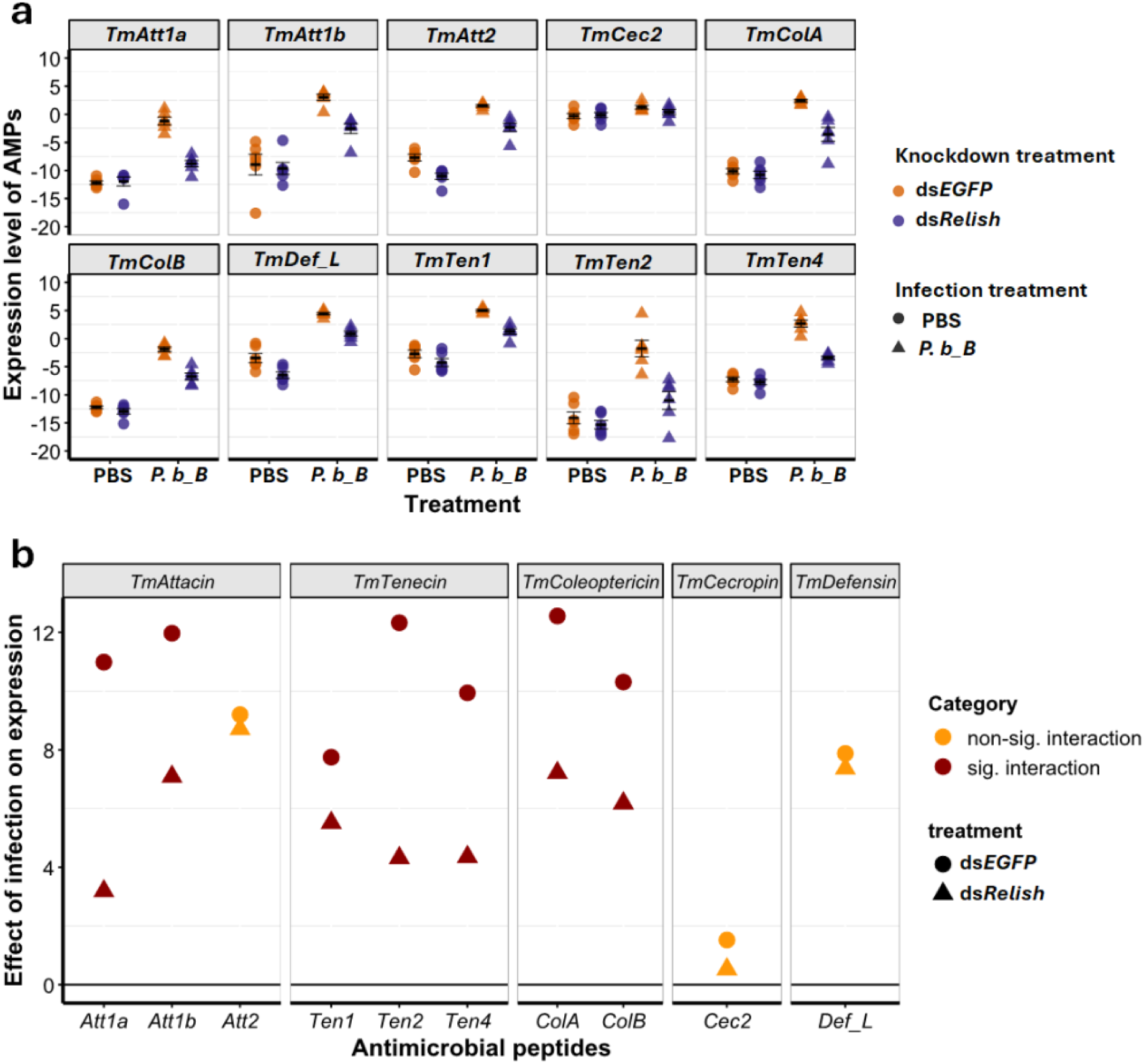
The effect of *Relish* knockdown on AMPs expression post-infection. (**a**) RT-qPCR analysis showing levels of *TmAtt1a*, *TmAtta1b*, *TmAtt2, TmTene1*, *TmTene2*, *TmTene4*, *TmColA*, *TmColB*, *TmCec2*, and *TmDef*-*L* mRNAs in whole-body extracts of ds*EGFP* (orange) and ds*TmRelish*-treated larvae (purple) infected with *P. burhodogranariea_B* (*P. b_B*, triangles) at 24 h post-infection. PBS-injected larvae (circles) were used as the mock control. *TmL27a* was used as the normalization control. Data points represent individual biological replicates, and error bars denote the mean ± SEM. (**b**) Estimates of linear models for changes in expression levels when comparing *P. b_B* infection with PBS control, in *TmRelish* knockdowns (triangles) or *EGFP* controls (circles). The horizontal line represents no change in AMPs expression. Points above indicate *P. b_B*-mediated upregulation of AMP expression in both *TmRelish* knockdowns and ds*EGFP* controls. Significance is color-coded: dark red (significant interaction between *TmRelish* knockdown and infection), and orange (no interaction).

We did not find a significant interaction between knockdown treatment and infection on expression levels of *TmAtt2*, *TmDef_L* and *TmCec2* (Fig. 4, Table S1). These AMPs were upregulated following infection without any apparent influence of the knockdown treatment on the magnitude of this upregulation. While for *TmCec2*, there was no statistically significant effect of knockdown treatment, for *TmAtt2* and *TmDef_L*, we found a significant downregulation in knockdown larvae (independently of infection) (Fig. 4, Table S1).

### The effect of pathogen infection on gut microbiota in tripartite interactions

When assessing the impact of *P. b_B* infection and Imd-mediated *AMP* expression on gut microbiota, we did not find a statistically significant interaction between both predictors (X² _1, 127_ = 0.02, *p* = 0.896) (Fig. 5, Fig. S5). Nevertheless, gut microbiota load was higher in infected individuals (infection: X² _1, 127_ = 4.87, *p* = 0.027), as well as in *TmRelish* knockdown individuals (ds*EGFP*/ds*TmRelish*: X² _1, 127_ = 20.30, *p* < 0.001) (Fig. 5, Fig. S2). We also found no indication for differences in the magnitudes between the effects of infection and knockdown treatments. *TmRelish* knockdown increased the microbiota load by a factor of ∼36 (95% CI: 7.60, 172.44), while it was ∼9.37 times higher for *P. b_B* infection (95% CI: 1.28, 68.35). Due to the widely overlapping confidence intervals we found no indication that microbiota load increased by different magnitudes following *TmRelish* knockdown and infection. Taken together these results suggest that while infections and immune system impairment impact the gut microbiota, we found no support for our prediction (P1) that pathogen infection affects gut microbiota and that this effect should be absent in *TmRelish*-knockdown larvae.

**Fig 5.**
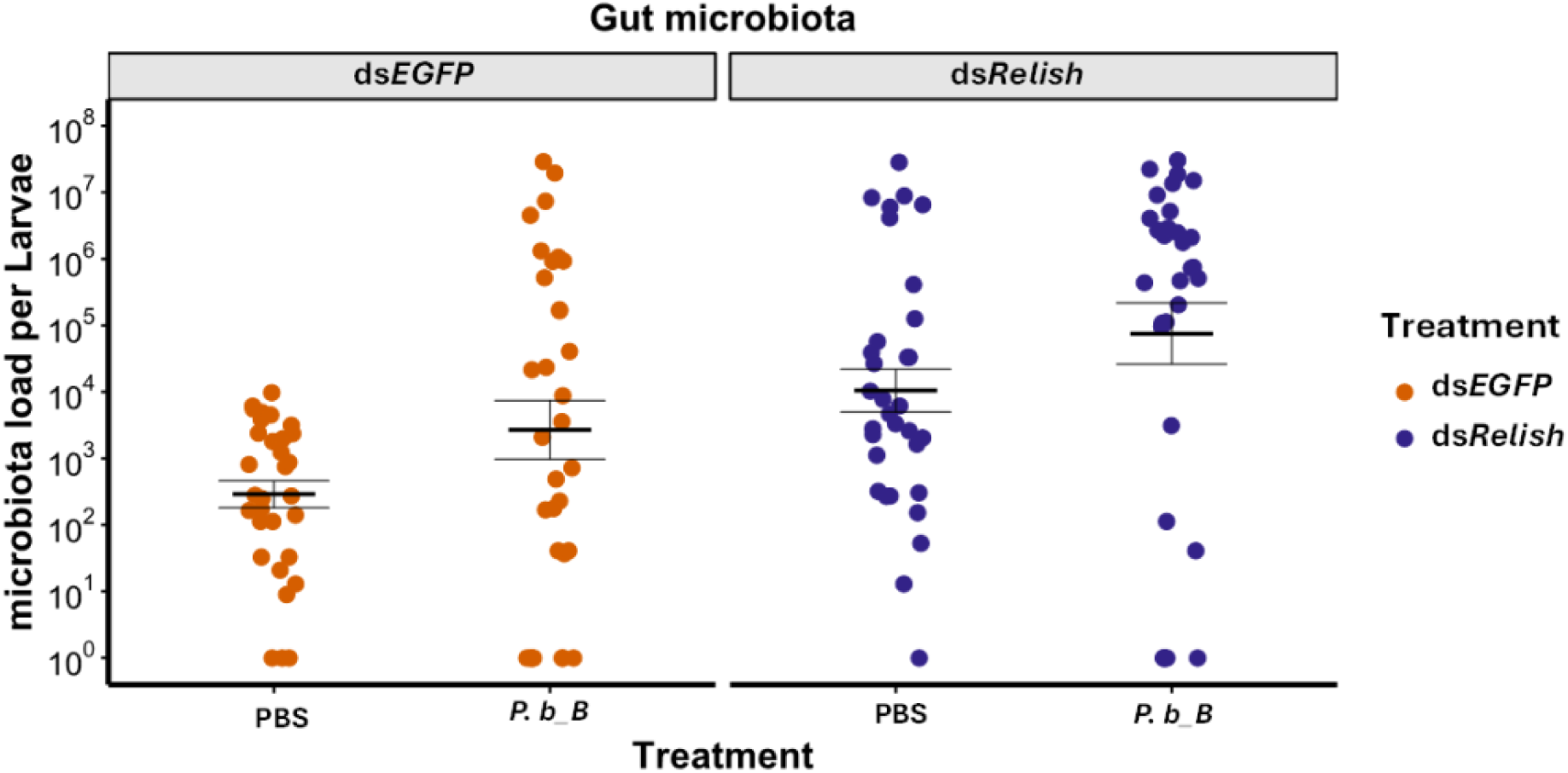
*TmRelish* knockdown and *Providencia burhodogranariea_B* infection effects on gut microbial abundance. Gut microbial load of all culturable bacteria in *Tenebrio molitor* larvae treated with ds*EGFP* and ds*TmRelish*, following exposure to either *P. burhodogranariea_B* (*P. bB*) or PBS. Means and standard errors are represented by the black lines, while the dots represent the biological replicates from individual guts (n=∼16 per treatment, two independent experiments). While we did not find a statistically significant interaction between both predictors (X² _1, 127_ = 0.02, *p* = 0.896), infection and *TmRelish* knockdown had a statistically significant effect (infection: X² _1, 127_ = 4.87, *p* = 0.027; ds*EGFP*/ds*TmRelish*: X² _1, 127_ = 20.30, *p* < 0.001).

### The effect of gut microbiota on pathogen infections in tripartite interactions

When investigating host survival in the presence or absence of gut microbiota following infection, we found no significant interaction between gut microbiota and knockdown treatments (X² _1, 32_ = 0.15, *p* = 0.70) (Fig. 6). However, we observed higher mortality on day 7 in AB-treated larvae compared to CR-treated larvae (X² _1, 32_= 4.63, *p* < 0.01), as well as ds*TmRelish*-treated larvae compared to ds*EGFP* larvae (X² _1, 32_ = 62.26, *p* < 0.001) (Fig. 6). These results suggest that both gut microbiota and immune system impairment impact the host survival, but do not support our prediction (P2) that gut microbiota influence pathogen infection and that this influence should be absent or reduced in *TmRelish*-knockdown larvae. Of note, to clarify the effects of antibiotics on our results, we ensured that after dsRNA treatment, larvae were fed the same diet but without chloramphenicol, preventing gut microbiota recovery. Additionally, we confirmed that *P. b_B* remained viable in the AB-treated group, suggesting no toxicity of antibiotic, as demonstrated in a separate experiment (data not shown).

**Fig 6.**
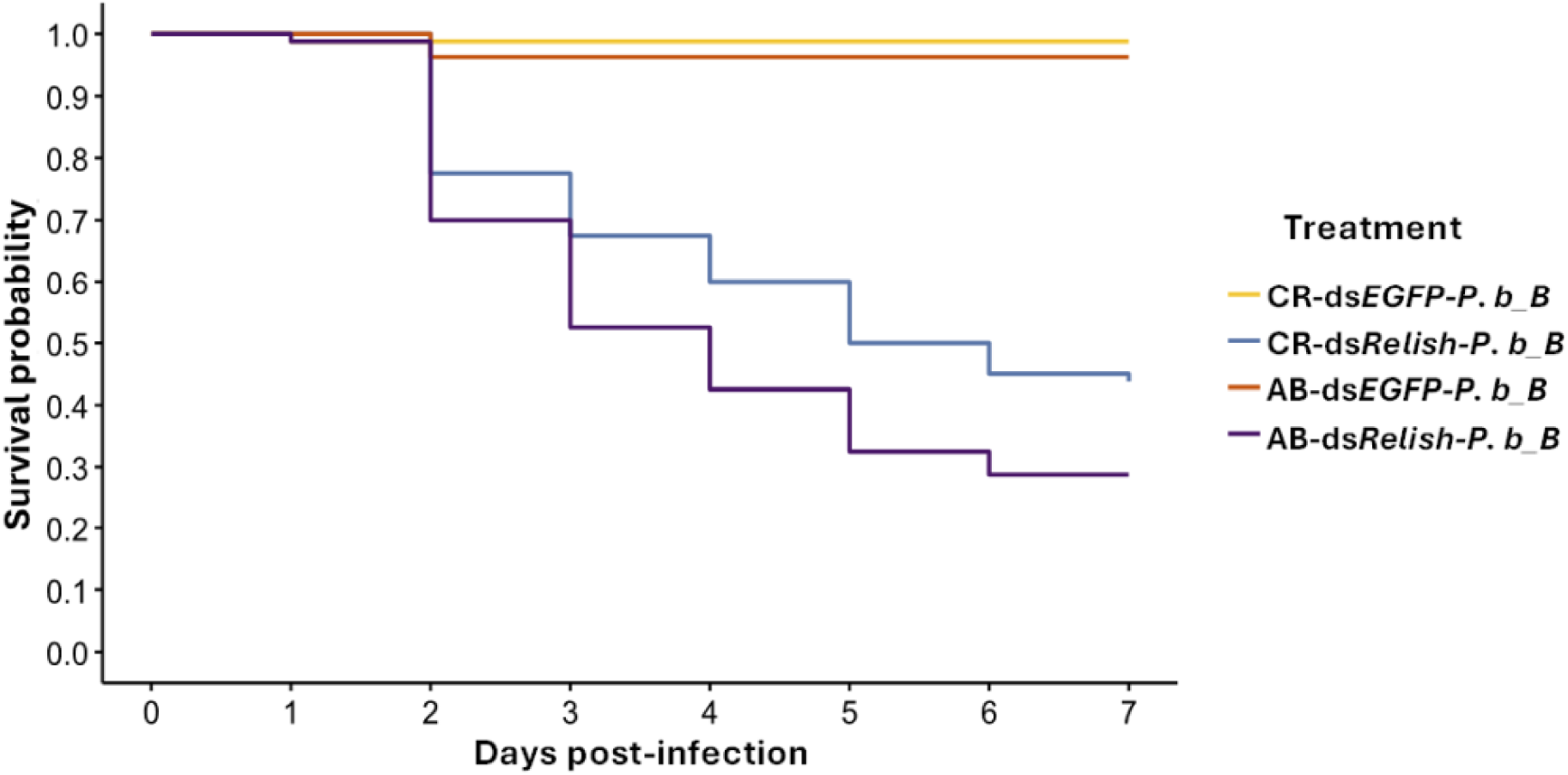
Effect of gut microbiota on pathogenicity of *Providencia burhodogranariea_B* in *TmRelish* knockdown larvae. Survival of control-treated or antibiotic-treated larvae injected with ds*EGFP* and ds*TmRelish* (n=∼10 per treatment, eight independent experiments) following *P. burhodogranariea_B* (*P. b_B*) infection was monitored over a period of seven days. While we did not find a statistically significant interaction between both predictors (X² _1, 32_ = 0.15, *p* = 0.70), gut microbiota and *TmRelish* knockdown had a statistically significant effect (CR/AB: X² _1, 32_= 4.63, *p* < 0.01; ds*EGFP*/ds*TmRelish*: X² _1, 32_ = 62.26, *p* < 0.001).

### AMP expression in tripartite interactions

The results above did not support our predictions, prompting us to investigate whether the presence of gut microbiota alters the dynamics of downstream genes that drive AMP synthesis and whether other pathways influence their expression. We found a statistically significant interaction between gut microbiota and infection on the expression levels of the AMPs *TmAtt1b*, *TmTen4*, *TmColB*, and *TmDef_L*, revealing that their expression is influenced by both the presence of gut microbiota and infection (Fig. 7, Table S2). Specially, while infections generally led to an upregulation of AMPs compared to non-infected PBS controls, for these four AMPs this upregulation was stronger in the presence of gut microbiota compared to AB- treated larvae (Fig. 7, Table S2). Thus, our analysis demonstrates that at least one AMP from each family, specifically *TmAtt1b*, *TmTen4*, *TmColB*, and *TmDef_L*, is more responsive to alterations in gut microbiota during infection compared to non-infection condition. In addition, for *TmAtt1a*, *TmTen1*, *TmTen2*, *TmColA*, and *TmCec2*, we found that infections lead to an upregulation independently of the presence of gut microbiota (Fig. 7, Table S2).

**Fig 7.**
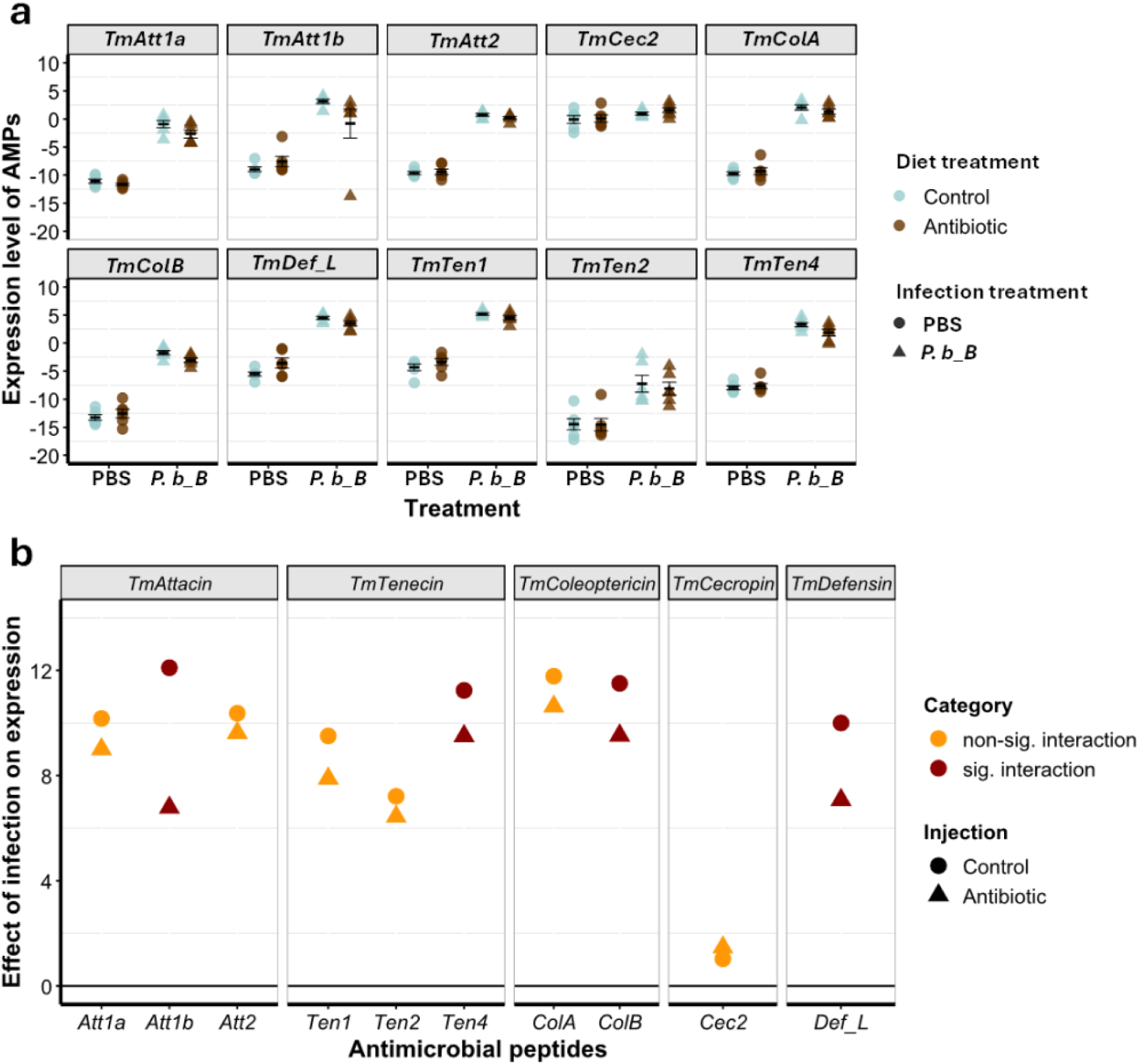
Regulation of antimicrobial peptide expression by gut microbiota in response to *Providencia burhodogranariea_B* infection. (**a**) RT-qPCR analysis of AMP gene expression including, *TmAtt1a*, *TmAtta1b*, *TmAtt2, TmTene1*, *TmTene2*, *TmTene4*, *TmColA*, *TmColB*, *TmCec2*, and *TmDef*-*L* in whole-body extracts of control (cyan) or antibiotic (brown) treated *Tenebrio molitor* larvae collected 24 h after infection with *P. burhodogranariea_B* (*P. b_B*, triangles). PBS-injected larvae (circles) were used as the mock control. Expression levels were normalized to *T. molitor 60S ribosomal protein L27a* (*TmL27a*) using delta Ct method (mean Ct of *TmL27a* – mean Ct of AMP gene). Data points represent individual biological replicates, each with two technical replicates (n = 6 larvae per treatment), and error bars denote the mean ± SEM. (**b**) Changes in expression levels of *TmAttacin* (*Att1a*, *Att1b*, *Att2*), *TmTenecin* (*Ten1*, *Ten2*, *Ten4*), *TmColeoptericin* (*ColA*, *ColB*), *TmCecropin* (*Cec2*), and *TmDefensin-like* (*Def_L*) were measured in control- (circles) and antibiotic-treated (triangles) larvae following infection with *P. burhodogranariea_B* or PBS control. The horizontal line represents no change in AMPs expression. Points above indicate *P. b_B*-driven upregulation of AMP expression. Significance is color-coded: dark red (significant interaction between gut microbiota and infection), and orange (no interaction).

## Discussion

In this study, we confirmed that bipartite interactions between the *Tenebrio* immune system and either gut microbiota or *P. burhodogranariea_B* infection are mediated through Imd-dependent AMPs. In tripartite interactions, we found no indication that indirect interactions between gut microbiota and pathogens are mediated by Imd-dependent AMPs. However, Imd-independent variation in AMPs appears to modulate the indirect interaction between gut microbiota and pathogens. Our findings align with the emerging paradigm of tripartite interactions (host-pathogen-microbiota) in shaping pathogen virulence evolution, suggesting that variation in host immunity can influence gut microbiota dynamics and impose selective pressures on the pathogen [1].

Our results, in line with observations in other insects, confirm an important role of the *Relish*- Imd pathway in regulating gut microbiota load and composition in *Tenebrio* larvae. The insect immune system, particularly Imd-mediated AMPs, has been implicated in shaping gut microbiota by selectively modulating bacterial populations, preserving homeostasis, and preventing dysbiosis [23, 37]. Consistent with *Drosophila* studies [21, 23], *TmRelish*- knockdown larvae exhibited an increase in gut microbial load and significant shifts in microbial composition. Here, we found that the genera *Bacillus* and *Pediococcus*, both Gram-positive bacteria with DAP-type peptidoglycan that activate the Imd pathway via peptidoglycan recognition proteins (PGRPs) [38, 39], were significantly reduced in ds*TmRelish*-treated larvae compared to controls. This suggests that *TmRelish* plays a key role in regulating their proliferation and maintaining gut stability (Fig. 2c-e).

*TmRelish*-knockdown significantly increased larval mortality following *P. b_B* infection, coinciding with enhanced bacterial survival in ds*TmRelish*-treated larvae (Fig 3). This susceptibility is likely due to the downregulation of systemic *AMP* families, including *TmAttacin* (except *TmAtt2*), *TmTenecin*, and *TmColeoptericin*. The expression of *TmAtt2*, *TmDef_L*, and *TmCec2* was not significantly affected, suggesting that these *AMP*s might be regulated by alternative immune pathways independent of *TmRelish* (Fig 4). While *TmCec2* is generally effective against Gram-negative bacteria such as *E. coli* [30, 40], its expression was not significantly induced in response to *P. b_B* infection. Of note, AMP expression in non-infected larvae remained largely unchanged (Fig. 4), underscoring the role of *TmRelish* in infection-specific immune modulation. These results align with previous studies showing that the Imd pathway recognizes pathogen-associated molecular patterns (PAMPs) via peptidoglycan recognition proteins (PGRPs), triggering AMP production [41]. In *T. molitor* and other insects, Imd-mediated AMPs play a crucial role in defending against Gram-negative bacteria.

Despite the well-established role of Imd-mediated AMPs in shaping bipartite interactions, contrary to our initial hypothesis, our results do not support their involvement in mediating indirect interactions between gut microbiota and *P. b_B* infection (Fig. 5, 6). While previous studies in tripartite contexts have highlighted the involvement of host immunity, such as the Toll pathway and Duox in mosquitoes [13, 17], we found no evidence that the Imd pathway serves a similar function in *T. molitor*. A possible explanation is that AMPs are also regulated by other pathways, which could, in turn, influence indirect interactions. In *TmRelish*- knockdown larvae, during infections AMPs were less strongly but still clearly upregulated (Fig. 4), suggesting that residual AMPs, potentially expressed through crosstalk with other immune pathways, might still have potential effects.

To investigate the possibility of Imd-independent AMPs expression, we examined AMPs expression in CR- and AB-treated larvae upon infection and non-infection controls. Our results suggest that Imd-independent AMPs expression indeed significantly modulates the indirect interaction between pathogen infection and gut microbiota (Fig. 7). When investigating AMP expression, we found a significant interaction between knockdown treatment and infection for at least one of AMPs from each family, including *TmAtt1b*, *TmTen4*, *TmColB*, and *TmDef_L*., This result suggests another immune pathway, independent of Imd, regulates these AMPs. This could be explained by distinct immune system architecture of *T. molitor* which differs from that of *Drosophila*. In *Drosophila*, AMPs are typically expressed in response to Gram-negative and Gram-positive bacteria. However, this specificity appears to be less pronounced in *T. molitor*, where *TmTen4* is induced by multiple ligands, including those activating both the Toll pathway (e.g., β-1,3-glucan and lysine-type peptidoglycan) and the Imd pathway (e.g., meso-DAP type peptidoglycan), suggesting a broader immune response [32]. It should also be noted that AMPs alone may be insufficient to mediate indirect effects in *Tenebrio*. Earlier studies in *Drosophila* have revealed the crosstalk between ROS and AMPs in regulating the indirect interactions between gut microbiota and pathogens. ROS, produced by Duox in response to pathogen infection, can regulate the Imd pathway and potentially AMPs activity, thereby altering gut microbiota load and composition [29]. Here, we focused only on the effects of Imd-mediated AMPs, rather than on the impact of other pathways like Duox-ROS. This may explain the lack of significant interactions between gut microbiota and pathogens in our study. Alternatively, it is possible that indirect interactions do occur but were not detected due to experimental limitations. For instance, the route of infection or time-dependent dynamics may influence these interactions, and if they only manifest at specific time points, our sampling may have missed these effects.

Our investigations of tripartite interactions also demonstrate a positive effect, between microbes, where *P. b_B* infection promotes gut microbiota load (Fig 5). This aligns with previous studies, such as fungal infection in *A. stephensi*, which altered immune responses in a way that promoted the gut microbiota growth [13]. Similarly, De Lorgeril et al. (2018) observed a comparable pattern in the oyster *C. gigas*, where viral infection suppressed immune responses, enabling microbiota to flourish. In both cases, as in our study, the pathogen indirectly promoted microbiota growth, reinforcing the idea that immune suppression or dysregulation can create favourable conditions for both microbiota and pathogens to coexist temporarily, until host mortality disrupts this balance [42]. However, in contrast to previous studies where microbiota proliferation increased host mortality, we observed lower host death with gut microbiota, suggesting a different host-microbiota-pathogen dynamic in *T. molitor* infected with *P. b_B*.

Our survival study is consistent with apparent competition, specifically a negative effect of microbiota on pathogens. Consistent with previous studies [43], AB-treated larvae exhibited higher mortality following *P. b_B* infection compared to CR-treated larvae (Fig. 6), indicating that the presence of gut microbiota provides protection against infection, likely by priming host immunity. Gut microbiota-pathogen interaction influencing infection outcomes have been observed in other insects. For example, *Aedes aegypti*, gut microbiota modulate dengue virus infection by stimulating the Toll pathway [17], Similarly, in *Anopheles gambiae*, gut microbiota limit *Plasmodium* infection by enhancing immune responses [18]. These studies involved oral infections, where gut microbes and pathogens interact directly within the midgut. However, apparent competition has also been observed following systemic infections. For instance, *Wolbachia*-infected *Drosophila* exhibit increased resistance to viral infections, indirectly suppressing viral success through immune priming [19].Accordingly, tripartite interactions shape infection outcomes and influence the evolutionary trajectories of both the host and the pathogen [44], potentially driving pathogen virulence in either direction, either facilitating or constraining it [1].

## Conclusion

Gut microbiota dynamic within host-pathogen interaction can either facilitate or prevent infections. Our findings provide insight into the indirect interactions between gut microbiota and pathogens, mediated through Imd-dependent and -independent AMPs. In summary, our model confirms bipartite interactions between *Tenebrio* Imd-dependent AMPs and either gut microbiota or *P. b_B* infection. However, in tripartite interactions, we found no evidence that indirect gut microbiota-pathogen interactions are mediated by Imd-dependent AMPs. Instead, these interactions appear to be regulated by Imd-independent AMPs. Host immunity thus plays a central role in shaping interactions with both gut microbiota and pathogens, maintaining microbial balance while defending against infections. By affecting virulence, these tripartite interactions are expected to impose selective pressures, which could influence the evolution of pathogen virulence.

## Material and methods

### Tenebrio molitor rearing

Late instar larvae (18^th^ to 20^th^ instar, approximately 2.5 - 3 cm in length) were obtained from a commercial supplier (Reptile Food Handels-u. Zucht GmbH, Berlin, Germany) and maintained in cohorts of 500 larvae in the dark at 25 ± 3°C and 60±5% relative humidity. The larvae were provided with an *ad libitum* supply of wheat bran, serving as their primary food source. For hydration and supplementary nutrition, a fresh apple slice was added to each container every 48 hours. At each feeding, the larvae were carefully checked, and newly pupated individuals were separated from the colony. Newly emerged adults were then transferred to separate containers for oviposition.

### Generation of antibiotic-treated larvae

*Tenebrio* larvae (8^th^ to 9^th^ instar, approximately 0.9- 1 cm in length) were fed on a control diet (CR) consisting of 100 g of wheat bran, 10 g soy protein, 20 g soy flour, 171 g of a 5% yeast and wheat flour mixture, and 0.2 mL propionic acid in 200 mL of distilled water for 5 days to establish similar gut microbiota composition. Whereas, early instar larvae (3^rd^ to 4^th^ instar, approximately 0.5 cm in length) were fed an antibiotic-treated diet (AB) consisting of 100 g of wheat bran, 10 g soy protein, 20 g soy flour, 170 g of a 5% yeast and wheat flour mixture, 0.5 g of chloramphenicol, 0.5 g of sorbic acid, 0.5mL of propionic acid in 200 mL of distilled water for 15 days. The last two days of feeding on the AB diet were limited to a diet with similar ingredients, except without chloramphenicol, to reduce the effect of the antibiotic. Fresh food was provided every 48 hours to avoid desiccation.

### Bacterial culture

*P. burhodogranariea_B* (provided by Sophie Armitage) was retrieved from a 50% glycerol stock and streaked onto Luria-Bertani (LB) agar plates [45]. Single colonies were inoculated into 100 mL of LB broth and cultivated under aerobic conditions at 30°C for 16 hours. Following incubation, 50 mL of the overnight culture was centrifuged at 2880 xg for 10 minutes at 4 °C. The resulting cell pellet was washed twice with 40 mL of phosphate buffered saline (PBS) and resuspended in 5 mL of PBS. Optical density at 600 nm (OD600) was measured using spectrophotometer (Ultrospec 10, Amersham Biosciences) across 1:10, 1:100, and 1:1000 dilutions. Then, bacterial concentrations were calculated based on the linear equation for *P. burhodogranariea_B* (y = 9 × 10⁸ (x) - 1 × 10⁷), allowing the desired colony-forming units per microliter (CFU/mL).

### Experimental procedures

All experimental individuals fed on either CR or AB were 9^th^ to 10^th^ instar larvae (approximately 1 - 1.5cm in length) [46].

Given that diet shapes gut microbiota, to determine if chloramphenicol in the diet affects gut microbiota composition, individual guts of CR- and AB-treated larvae (n=12 per treatment) was dissected and bead-grounded for DNA extraction. Differences between treatments were assessed using both culture-dependent and culture-independent approaches. Detailed procedures are provided in the supporting information (SI Appendix, Fig. S3).

To examine potential impacts of the NF-κB transcription factor, Relish-Imd, on gut microbiota load and composition, an RNAi knockdown construct targeted against *TmRelish* was expressed, accordingly, conventional larvae were treated with double-stranded RNA (dsRNA) of *Enhanced Green Fluorescent Protein* (ds*EGFP*) and ds*TmRelish*. We recapitulated our previous finding on knockdown efficiency, result was confirmed via qRT-PCR [30] (SI Appendix, Fig. S4). Guts from both groups (n=∼12 per treatment, six biological replicates) were dissected in PBS at 3-day post-injection. Detailed procedures are provided in the supporting information (SI Appendix).

To investigate whether *P. b_B* infection, *TmRelish* knockdown, and their interaction impact the host survival and *Relish*-dependant AMPs expression, conventional larvae were first injected with ds*EGFP* or ds*TmRelish*. Three days post-dsRNA exposure, larvae were injected with 2µL of PBS (control) or with *P. burhodogranariea_B* (*P. b_B*) at a concentration of 5 × 10^7^ colony forming units (CFU/mL), corresponding to a dose of approximately 10^5^ CFU per insect. The bacterial dose was selected as the highest dose that did not cause mortality (SI Appendix, Fig. S5). For host survival analysis, each group of insects, including control (*P. b_B* without dsRNA injection, CR-*P. b_B*), ds*EGFP*-*P. b_B*, ds*TmRelish*-*P. b_B* (n=∼30 per treatment, three independent experiments) was monitored daily for one week. Additionally, we assessed *P. b_B* colonization in RNAi-treated larvae. To address bacterial survival, ds*EGFP*-*P. b_B* and ds*TmRelish*-*P. b_B* (n=∼10 per treatment per day) were bead-grounded at different time points 0-day, 1-day, 2 days, 3 days, 4 days, 5 days, 6 days, and 7 days post-infections. The data represent at least two independent experiments. For measuring the transcript level of targeted mRNA, i.e., *TmRelish*-mediated AMPs expression, six larvae as a biological replicate were collected from larvae treated with ds*EGFP* and ds*TmRelish* followed infection with either PBS or *P. burhodogranariea_B* (ds*EGFP*-PBS, ds*TmRelish*-PBS, ds*EGFP*-*P. b_B*, ds*TmRelish*-*P. b_B*).

Next, to investigate whether pathogen infection affects the gut microbiota and further assess whether this effect differs in *TmRelish* knockdown individuals (Fig 1a), conventional larvae treated with ds*EGFP* and ds*TmRelish* followed infection with either PBS or *P. burhodogranariea_B* (ds*EGFP*-PBS, ds*TmRelish*-PBS, ds*EGFP*-*P. b_B*, ds*TmRelish*-*P. b_B*) were sacrificed at 24 h post-infection and assessed for bacterial load (n=∼16 per treatment, two independent experiments).

To test knockdown of *TmRelish* effects on the host survival in the presence and absence of gut microbiota during infection (Fig 1b), CR- and AB-treated larvae were injected with ds*EGFP* or ds*TmRelish* followed infection with *P. b_B*. For host survival analysis, each group of insects, including CR-ds*EGFP*-*P. b_B*, CR-ds*TmRelish*-*P. b_B*, AB-ds*EGFP*-*P. b_B*, AB-ds*TmRelish*-*P. b_B* (n=∼10 per treatment, eight independent experiments) was monitored daily for one week. Additionally, since gut microbiota act as a regulator in maintaining a healthy gut and effective immune response through the constitutive expression of Imd-dependent AMPs [47, 48], we used RT-qPCR to investigate the effects of imbalanced gut microbiota on AMPs expression. For this, we injected CR- and AB-treated larvae with PBS or with *P. b_B*. Two independent experimental replicates were conducted. In each experimental replicate and for each treatment, three larvae were injected, giving a total of 24 larvae across the four groups CR-PBS, AB-PBS, CR-*P. b_B*, AB-*P. b_B*.

Of note, larvae were not surface sterilized prior to homogenization, in line with the recommendations by Hammer et al. (2015), suggesting that surface sterilization should be avoided in insect microbiota research, as it may inadvertently alter the composition of internal bacterial communities [49].

### DNA extraction and bacterial 16S sequencing analysis

DNA was extracted from homogenized larval gut tissue using DNeasy PowerSoil Pro Kit (Qiagen, Germany) with the following modifications. Larval guts were collected in 2 mL microcentrifuge tubes (Safe-Lock tubes, Eppendorf) containing 800 µL of CD1 solution buffer. After addition of two stainless steel beads (Ø 3 mm, Retsch), samples were homogenised using a tissue homogenizer (Mill MM400, Retsch) at 25 Hz for 10min. The resulting homogenates were incubated overnight with 10 µL of proteinase K at 56°C for 16 hours in a ThermoMixer^®^ C. The remaining steps were carried out according to the manufacturer’s protocol. DNA concentration was then measured via Qubit™ (Invitrogen) and subsequently diluted to 5 ng/µL.

The V2-V3 region of the 16S rRNA encoding gene was amplified using 10 ng DNA (or negative control) as template, 1.25 µL of 10µM primers 515 forward (515F) and 806 reverse (806R) (Table S3) and 12.5 µL of Q5 High Fidelity Mastermix (New England Biolabs) in a 25 µL reaction. The PCR amplicons were purified using CleanNGS CNGS-0050 (GC Biotech B.V.). Libraries were sequenced using an Illumina MiSeq (Illumina) using sequencing kit v3 600 cycles at the Berlin Center for Genomics in Biodiversity Research (BeGenDiv). Detailed procedures are explained in supporting information (SI Appendix).

The primer sequences were removed from the raw reads using cutadapt [50] and their quality checked with FastQC [51]. Accordingly, the raw reads were further trimmed at 200 bp for both forward and reverse reads with the ‘filterAndTrim’ function of the package ‘dada2’ [52]. They were then processed into amplicon sequence variants (ASVs) using Divisive Amplicon Denoising Algorithm 2 (DADA2) pipeline in R (‘dada2’ package). Taxonomy was assigned to the ASVs with the ‘assignTaxonomy’ function of the ‘dada2’ package, which uses a naive Bayesian classifier method of [53], and by querying against the Ribosomal Database Project (RDP) reference database [54]. Reads identified as mitochondria or chloroplasts, commonly observed in insect gut samples, were excluded from further analysis [55]. Bray-Curtis’s dissimilarity was then used to assess beta diversity, and Principal Coordinate Analysis (PCoA) was performed to visualize these differences in community structure.

### RT-qPCR

Total RNA was isolated by homogenizing larvae using TRIzol reagent following the manufacturer’s instructions (Direct-zol RNA Miniprep Plus Kits, ZYMO Research, Europe GmbH). RNA purity was validated via NanoDrop™ 2000/2000c Spectrophotometers (Thermo Scientific, Wilmington, DE, USA). The resulting mRNA was stored at -80°C until being used for real time qPCR. To set up the RT-qPCR for different genes, an amount of 20 ng/µL of the extracted mRNA was used using Power SYBR™ Green RNA-to-CT™ 1-Step Kit (Applied Biosystems ^TM^) with forward and reverse primers in a 10 µL reaction as previously described [30]. Briefly, forward and reverse primers were mixed with Power SYBR™ Green RT-PCR mix, RT-enzyme mix and nuclease-free water to amplify the following gene expression: *TmTenecin-1*, *TmTenecin-2*, *TmTenecin*-*4*, *TmAttacin-1a*, *TmAttacin-1b*, *TmAttacin-2, TmDefensin*-*like*, *TmColeoptericin-A*, *TmColeoptericin-B*, *TmCecropin,* and *T. molitor 60S ribosomal protein L27a* (*TmL27a*) (Table S3). The mean Ct value of the AMP gene of interest (GOI) was normalized using the mean Ct value of housekeeping gene by calculating delta Ct (mean Ct of housekeeping gene – mean Ct of GOI), where the housekeeping gene is *T. molitor 60S ribosomal protein L27a* (*TmL27a*) [56].

### cDNA synthesis and generation of double-stranded RNA

To synthesize PCR-based double-stranded RNA targeting *T. molitor* Relish (GenBank accession No. EEZ97717.1), a cDNA fragment corresponding to the target gene was produced using RevertAid™ Premium First Strand-cDNA-Synthese kit (Invitrogen), with total RNA extracted from the late instar larvae exhibiting high expression as template (Direct-zol RNA Miniprep Plus Kits, ZYMO Research, Europe GmbH). Gene-specific primers containing a T7 polymerase promotor sequence (TAATACGACTCACTATAGGG) at the 5’ end (SI Appendix, Table S5) were used to amplify the 851 bp PCR product from the cDNA template (KAPA2G Fast ReadyMix PCR Kit, KAPA Biosystems). PCR cycling profile were as follow: 95 °C for 2 min, followed by 30 cycles of denaturation at 95 °C for 20 s, annealing at 56 °C for 30 s, and extension at 72 °C for 5 min. Similarly, EGFP target (508 bp), cloned into the plasmid (pGEM T-easy-GFP, Promega) was amplified using gene-specific primers (Table S3). Following checking of amplicon size via 2% agarose gel, the PCR products were purified using the PCR/DNA Clean-Up Kit (Roboklon). The purified amplicons served as templates for *in vitro* transcription, using the HighYield T7 RNA Synthesis Kit (Jena Bioscience) in accordance with the manufacturer’s instructions. Next, the synthesized dsRNA was washed, and the resulting pellet was resuspended in nuclease-free water and stored at -20°C for future applications.

To assess the knockdown efficiency of *TmRelish*, conventional larvae were injected with 1000±100 ng of dsRNA (1000 μg/μL in 1 μL) of both ds*TmRelish* and ds*EGFP* (negative control). Total RNA was extracted from a pool of four larvae per day (n=4, three experimental replicates) at indicated time points of days 1, 3, 5, and 7 post-injection. RT-qPCR was performed as previously described protocol.

### Cultivation and quantification of gut microbiota

To assess the impact of *TmRelish* knockdown on the proliferation of *T. molitor* gut microbiota, each larva subjected to ds*TmRelish* or ds*EGFP* (n=15 per treatment) were dissected at three days post-exposure. Individuals were homogenised in 200 µL PBS using two sterile glass beads at 30 Hz for 20 seconds using a tissue homogenizer (Mill MM400, Retsch). Homogenates were serially diluted (1:10 to 1:10^5^) and plated onto the Trypticase soy agar (TSA) plates [57].

For examining the effects of *TmRelish* knockdown during infection on the gut microbiota, individual guts of ds*EGFP*-PBS, ds*TmRelish*-PBS, ds*EGFP*-*P. b_B*, ds*TmRelish*-*P. b_B* were dissected and plated onto either TSA or TSA medium containing 16 mg/mL of nalidixic acid and 32 mg/mL of rifampicin. After 16h of incubation at 37°C, CFUs were counted from five (5 μL) drops per sample. Each individual served as a biological replicate and each treatment contained at least three replicates.

### Bacterial survival

To examine if *P. burhodogranariea_B* can evade the impaired immune system of *T. molitor* or is eliminated by it, larvae treated with ds*EGFP*-*P. b_B* and ds*TmRelish*-*P. b_B* were homogenised in 250 μL of LB broth at different time points (0-day, 1-day, 2-day, 3-day, 4-day, 5-day, 6-day, and 7-day post-infections). Homogenization was performed as described above, followed by centrifugation at 420 xg for 1 minute at 4 °C. Homogenates from each individual were added into 180 μL of LB broth and then serially diluted (1:10 to 1:10^5^). *P. burhodogranariea_B* colonies were cultured on LB medium containing 16 mg/mL of nalidixic acid and 32 mg/mL of rifampicin for 16 h at 30°C. Five drops (5 μL) per larvae were counted and averaged as replicates. In addition, we injected PBS in ds*EGFP* and ds*TmRelish* larvae as a negative control: no colonies were recovered from these larvae (n =8).

### Statistical analysis

All data were analysed using R statistical software (v 4.4.1; R core team 2024) [58]. We fitted generalized linear mixed models (GLMMs), implemented in the “glmmTMB” package (version 1.1.9) [59], for the knockdown efficiency analysis, AMP expression analysis, survival assays, and gut microbiota load. When necessary, to better understand the effects of significant interactions we performed pairwise post hoc tests using “emmeans” package (package version 1.10.4) [60]. Assessment of model assumptions were evaluated using the DHARMa package (version 0.4.6) [61].

In bipartite interaction between host and gut microbiota, for analysis of knockdown efficiency of gene expression, we evaluated the effects of knockdown treatment on delta Ct values. Here, delta Ct was defined as the mean Ct of housekeeping gene minus mean Ct of gene of interest, with delta Ct serving as response variable and replicates as a random effect. For gut microbial load, we fitted a GLMM with a negative binomial distribution, to assess the effect of *TmRelish* knockdown on gut microbial load. The response variable was gut microbial load (CFUs), with knockdown treatments (ds*EGFP* or ds*TmRelish*) as the fixed effect and experimenter and replicate as random effects. The amplicon sequence variant (ASV) abundance data from the 16S sequencing of the larval bacterial gut community generated by the DADA2 pipeline, was imported and analysed with the ‘phyloseq’ package [62]. Reads identified as mitochondria, chloroplasts, or eukaryote plant material, commonly observed in insect gut samples, were excluded from further analysis [55]. Briefly, ASVs were agglomerated at the genus and order levels, and the data from the two experiments were subsetted and analysed separately. Operational Taxonomic Units (OTUs) counts for each sample were normalized to the total number of OTUs per sample, and a Bray-Curtis dissimilarities matrix was built as an index of beta diversity. A PERMANOVA was performed with the ‘adonis’ function of the ‘vegan’ package with an alpha level of 0.05 as a threshold to determine significance [63]. The phyloseq data was then converted to a DESeq2 dataset, and the geometric mean, size factors and dispersions of each OTU were estimated before performing a Likelihood Ratio Test (LRT) to identify differentially abundant OTUs (alpha = 0.05) between treatments using ‘DESeq2’ package [64].

In bipartite interaction between host and pathogen, for survivorship data were performed using a GLMM, with mortality at day 7, defined as a binary outcome (number of dead or alive) as response variable, knockdown treatment (ds*EGFP* or ds*TmRelish*) as fixed effect and date as a random effect to account for variability across days. To test the effects of knockdown treatment and time on log-transformed bacterial load as the response variable, we fitted a linear model, with knockdown treatment and time serving as fixed effect and replicates as a random effect. For AMP expression analysis, we fitted a GLMM to assess the effects knockdown treatment, infection, and their interaction as fixed effects on delta Ct values.

In tripartite interaction context, to evaluate the effects of knockdown treatment, infection, and their interaction on log-transformed gut microbiota load, we fitted a linear mixed model, with knockdown treatment and injection serving as fixed effects, gut microbiota load as response variable and replicates as a random effect. We analysed the data from survival and AMP expression similar to bipartite interaction experiments. For survivorship data we used a GLMM, with mortality at day 7 as response variable, knockdown treatment (ds*EGFP* or ds*TmRelish*) and effects of gut microbiota (CR- or AB-treatments) as fixed effect and date as a random effect. For AMP expression analysis, we fitted a GLMM to assess the effects of gut microbiota, infection, and their interaction as fixed effects on delta Ct values.

## Supplementary Information

The online version contains supplementary materials.

## Acknowledgements

We thank Sophie Armitage for gifting the bacteria strain.

## Authors’ contributions

MK conceived the project, designed the experiments, and wrote the manuscript. MK and HX conducted the experiments. SM and SS performed 16S rRNA sequencing, and CZ analysed the sequencing data. MK and MF analysed the data. MF and JR commented on the manuscript. JR procured reagents and materials.

## Data availability

The raw data set and R scripts used in all analyses are available from the corresponding author upon request.

## Declarations

### Ethics approval and consent to participate

Not applicable.

