## Supplementary material for "Immune-mediated indirect interaction between gut microbiota and bacterial pathogens"

**This file includes:**

**Supporting Information**

**Figure S1 to S6**

**Tables S1 to S4**

### Supporting Information

### SI Materials and Methods

#### Bacterial 16S sequencing

The V2-V3 region of the 16S rRNA gene was amplified under the following conditions: 98°C for 30s, 30 cycles of 98°C, touch down 63°-53°C (-1K every second cycle) 30s, 72°C 10s; and a final extension step 72°C 120s. The PCR amplicons were purified using CleanNGS and indexed with unique indexing primer combinations: 10µl purified PCR1, 1.25µl 10µM indexing primer P5, 1.25µl indexing primer P7, 12.5µl Q5 High Fidelity Mastermix (New England Biolabs) with these reaction conditions: 98°C 30s; 8 cycles of 98°C 10s, 67°C 30s, 72°C 30s; final extension 72°C 120s.

#### Knockdown efficiency of RNAi-treated in *Tenebrio molitor* larvae

We used RT-qPCR to evaluate the knockdown efficiency of ds*TmRelish* compared to ds*EGFP* in pooled samples of three larvae at 1-day, 3 days, 5 days, 7 days post-injection, and in individual larvae at 3 days post-injection. The Ct values were assessed using Power SYBR™ Green RNA-to-CT™ 1-Step Kit (Applied Biosystems TM) under PCR cycling profile as follow: 95 °C for 5 min (holding stage), followed by 40 cycles at 95 °C for 15 s (denaturation) and 60 °C for 30 s (annealing).

**SI Results**

**Effect of chloramphenicol on gut microbiota diversity in *Tenebrio* larvae**

We investigated whether disruption of bidirectional interaction via antibiotic-treated diet will affect the gut microbiota balance. Using both culture-dependent and culture-independent approaches, we assessed the gut microbiota composition of *Tenebrio* larvae in either conventionally reared (CR) or antibiotic-treated (AB). The results showed that AB-treated larvae had no culturable microbiome compared to CR-treated larvae (data not shown). Beta-diversity analysis revealed significant clustering patterns of microbial communities between AB and CR ($R^2 = 0.21$, $F = 5.98$, $p < 0.001$), suggesting a high degree of similarity in microbial communities within the AB group, and greater variability within the CR group (Fig. S2). Analysis showed that one taxon, from the genus *Bacillus* (family *Bacillaceae*), was significantly more abundant in the CR compared to the AB group (adjusted $p < 0.05$) (Fig. S2 b). Examination of 16S rRNA gene amplicon sequencing revealed that CR larvae had an increased relative abundance of operational taxonomic units (OTUs) predominantly belonging to the orders *Lactobacillales* (genus *Lactococcus*) and *Bacillales* (genus *Bacillus*), whereas AB larvae were more dominated by orders such as *Enterobacterales*, *Rhodospirillales*, *Rhizobiales*, and *Pseudomonadales* (Fig. S2 c-d). Taken together, these results suggest that CR group maintained a healthy and stable gut microbiota communities, while AB group not only exhibited a significant shift but also developed a distinct gut microbiota that is either resistant to chloramphenicol or is opportunistic colonizers in a disrupted microbiome.

***Tenebrio molitor* Relish knockdown efficiency**

We recapitulated our previous finding, showing that *TmRelish* is efficiently knocked down compared to ds*EGFP* on the third day post-dsRNA injection (ds*EGFP*/ds*TmRelish*: $t_{(31)} = 4.817$, $p < 0.001$) (Fig. S3 a). The respective knockdown remained significantly downregulated relative to the control at seven days post-injection (ds*EGFP*/ds*TmRelish*: $t_{(31)} = 3.939$, $p = 0.0004$) as measured by qRT-PCR (Fig. S3 a). *TmRelish* is efficiently knocked down compared to ds*EGFP* on the third day post-dsRNA injection in individuals (ds*EGFP*/ds*TmRelish*: $t_{(10)} = -6.861$, $p < 0.001$) (Fig. S3 b).

### Supplementary Figures

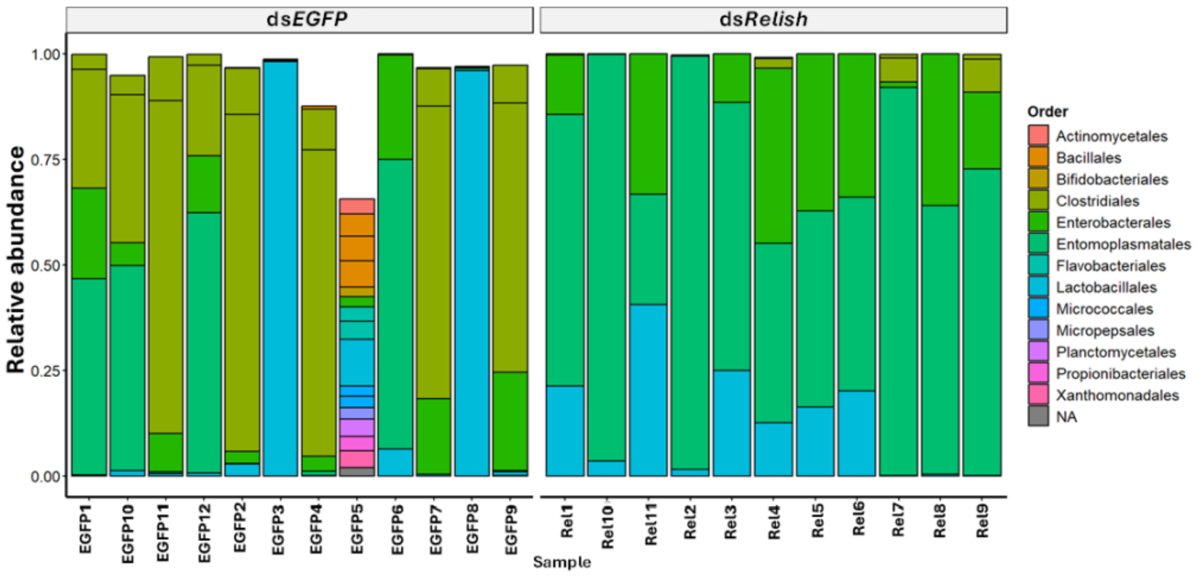


**Figure S1. Relative abundance and taxonomic assignment of the top 20 orders.** 16S rRNA gene sequencing in *T. molitor* larvae treated with ds*EGFP* and ds*TmRelish*, visualized by bar plots. Each bar represents an individual insect, with coloured box indicating different taxa. The hight of each box represents to the relative abundance of that taxon within the samples. Grey boxes indicate OTUs for which no taxonomy could be assigned.

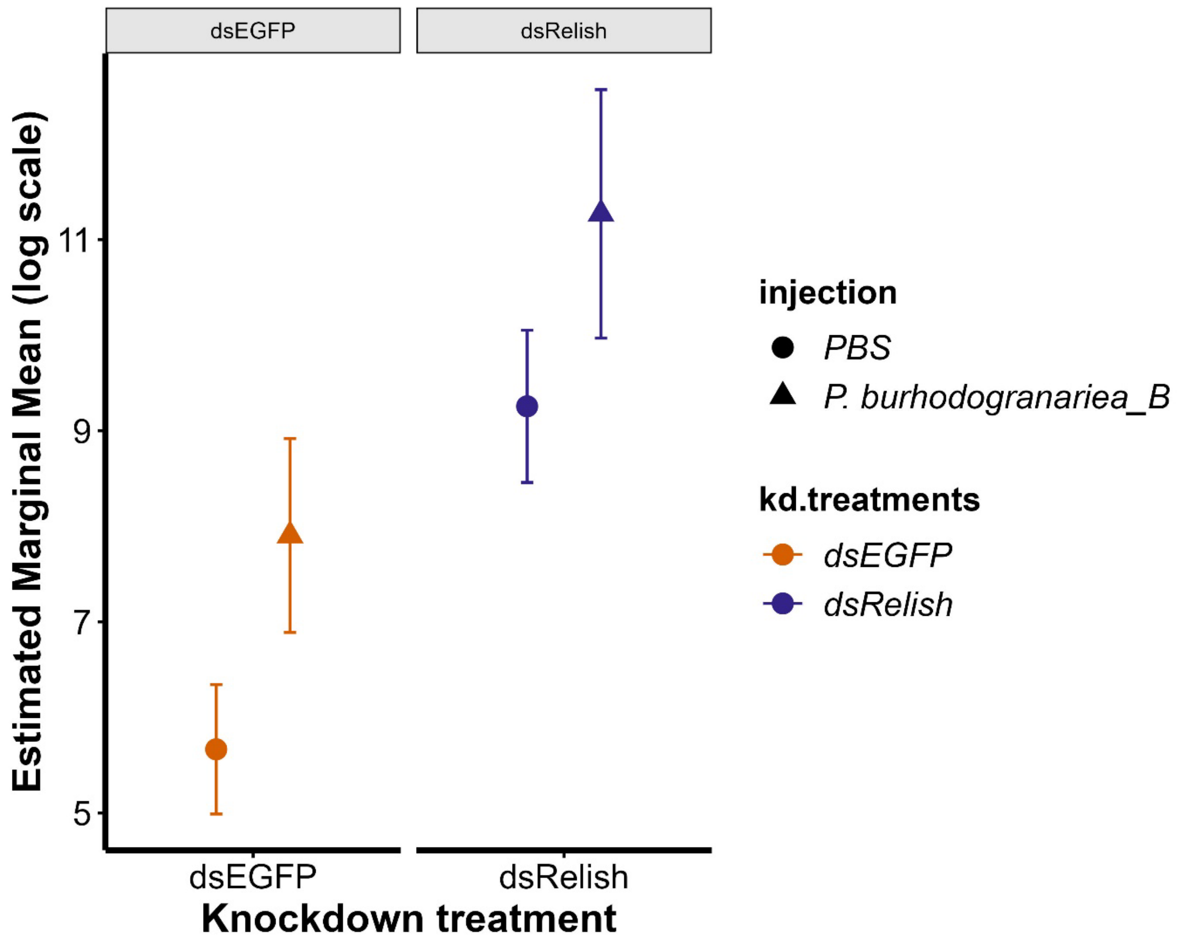


**Figure S2. Relish knockdown and *Providencia burhodogranariea_B* infection effect on gut microbiota.** Estimated marginal means with 95% confidence intervals of gut microbial load in *Tenebrio molitor* larvae treated with ds*EGFP* (orange) and ds*TmRelish* (purple), following exposure to either *P. burhodogranariea_B* (triangles) or PBS (circles).

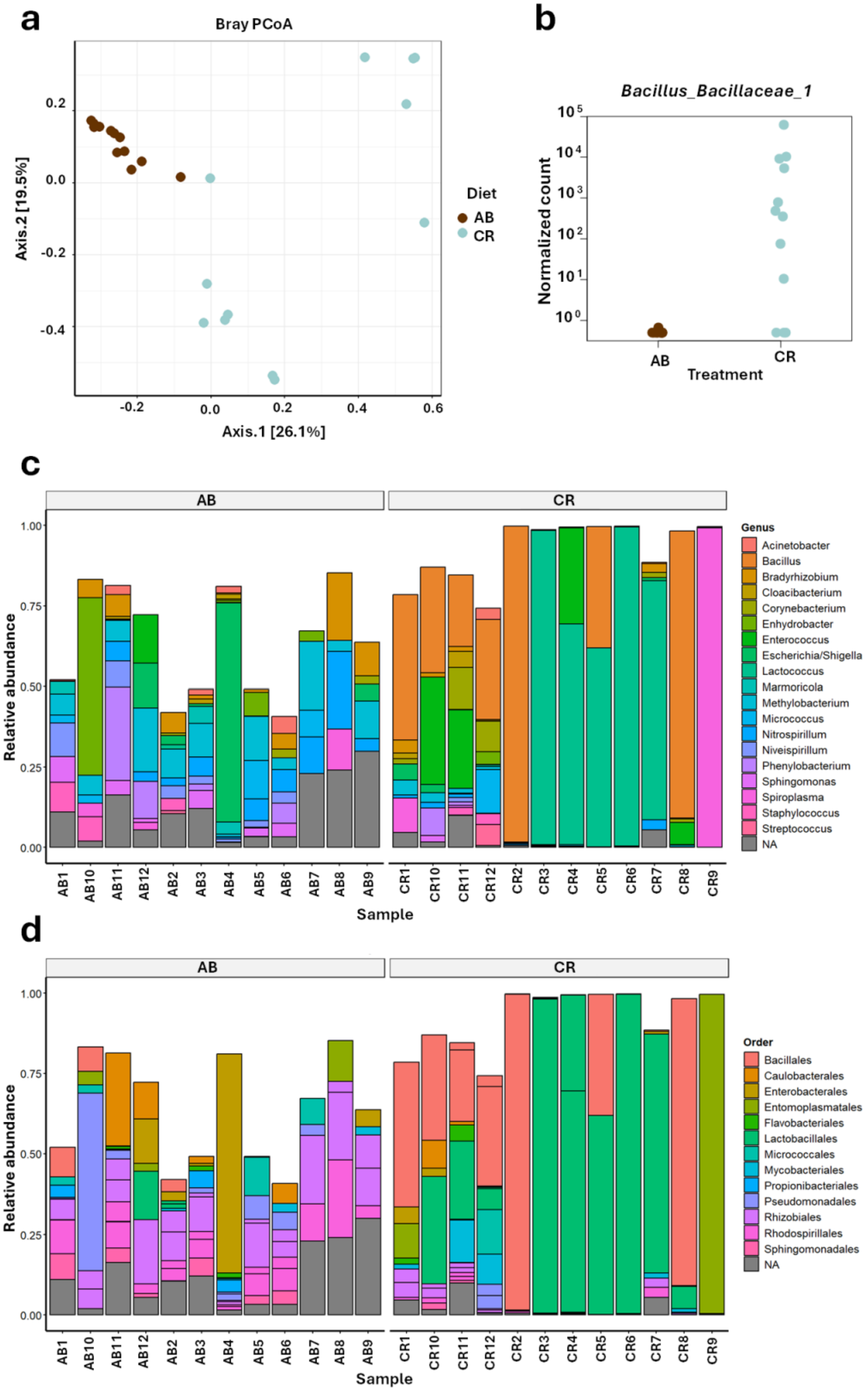

a
Bray PCoA
Axis.2 [19.5%]
Axis.1 [26.1%]
0.2
0.0
-0.2
-0.4
-0.2
0.0
0.2
0.4
0.6
Diet
AB
CR
b
Bacillus_Bacillaceae_1
Normalized count
$10^5$
$10^4$
$10^3$
$10^2$
$10^1$
$10^0$
AB
CR
Treatment
c
AB
CR
Relative abundance
1.00
0.75
0.50
0.25
0.00
Genus
Acinetobacter
Bacillus
Bradyrhizobium
Cloacibacterium
Corynebacterium
Enhydrobacter
Enterococcus
Escherichia/Shigella
Lactococcus
Marmoricola
Methylobacterium
Micrococcus
Nitrospirillum
Niveispirillum
Phenylobacterium
Sphingomonas
Spiroplasma
Staphylococcus
Streptococcus
NA
AB1
AB10
AB11
AB12
AB2
AB3
AB4
AB5
AB6
AB7
AB8
AB9
CR1
CR10
CR11
CR12
CR2
CR3
CR4
CR5
CR6
CR7
CR8
CR9
Sample
d
AB
CR
Relative abundance
1.00
0.75
0.50
0.25
0.00
Order
Bacillales
Caulobacterales
Enterobacterales
Entomoplasmatales
Flavobacteriales
Lactobacillales
Micrococcales
Mycobacteriales
Propionibacteriales
Pseudomonadales
Rhizobiales
Rhodospirillales
Sphingomonadales
NA
AB1
AB10
AB11
AB12
AB2
AB3
AB4
AB5
AB6
AB7
AB8
AB9
CR1
CR10
CR11
CR12
CR2
CR3
CR4
CR5
CR6
CR7
CR8
CR9
Sample

**Figure S3. Microbiota composition and differential abundance in control and antibiotic-treated *Tenebrio molitor* larvae.** (**a**) Principal coordinates analysis (PcoA) plot visualizing Bray-Curtis dissimilarity in the composition of the bacterial gut microbiota between control (cyan) or antibiotic-treated (brown) *Tenebrio molitor* larvae ($9^{th}$ to $10^{th}$ instar). Each point represents the Bray-Curtis index of an individual larva. The axis labels indicate the percentage of variation captured by each dimension. A PERMANOVA with 999 permutations showed that the treatment significantly separates the samples. (**b**) ASV counts of an OTU of the *Bacillus* genus (family *Bacillaceae*) whose abundance is significantly explained by the treatment (CR or AB). Each point represents the ASV count in a single *T. molitor* larva of the control (CR, cyan) and antibiotic-treated (AB, brown) groups. (**c**) Relative abundance of genera (**d**) and orders for the top 20 taxa detected by 16S rRNA gene sequencing in *T. molitor* larvae treated with different diets, visualized by bar plots. Each bar represents an individual sample, with coloured box indicating different taxa. The hight of each box represents the relative abundance of that taxon within the samples. Grey boxes indicate OTUs for which no taxonomy could be assigned.

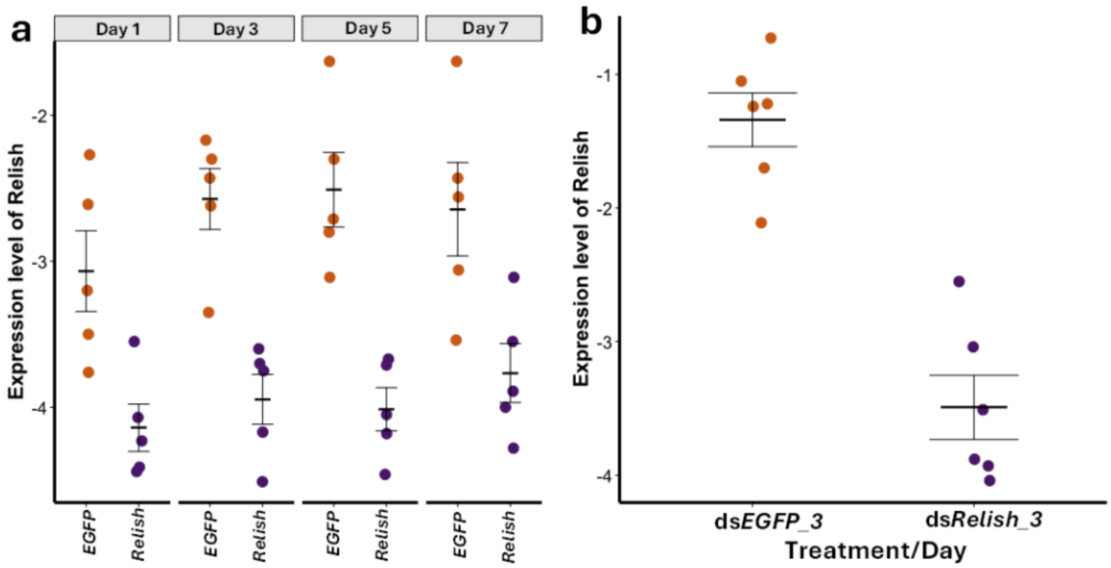


**Figure S4. Knockdown efficiency of *TmRelish* in RNAi-treated *Tenebrio molitor* larvae.** Quantitative measurement of *Tenebrio molitor Relish* (*TmRelish*) mRNA level in ds*TmRelish*-injected larvae determined by RT-qPCR. One microgram of dsRNA targeting *TmRelish* at the concentration of 1000 ng/µL in 1 µL were injected to $9^{th}$ to $10^{th}$ instar larvae. Total RNA was isolated on the $1^{st}$, $3^{rd}$, $5^{th}$, and $7^{th}$ day following treatments (n = 3 pools of three beetles each per treatment group per time point). (**a**) Knockdown efficiency of *TmRelish* in individuals was measured on the third day post-exposure. (**b**) The mRNA quantity of *TmRelish* was measured in relation to *T. molitor 60S ribosomal protein L27a* (*TmL27a*) as an internal control. EGFP RNAi was used as a negative control. Dots indicate the data. The Ct values of the gene of interest were normalized to the Ct values of the reference gene.

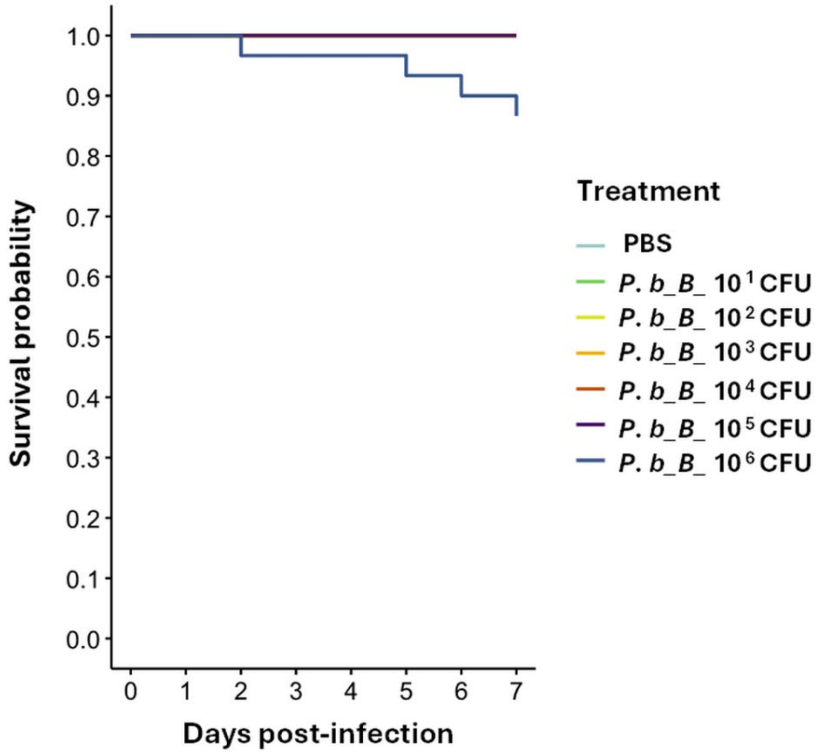


**Figure S5. Survival of *Tenebrio molitor* larvae upon infection with different doses of *P. burhodogranariea_B*.** Conventional *T. molitor* ($9^{th}$ to $10^{th}$ instar) larvae were infected with *P. burhodogranariea_B* at doses of $10^1$, $10^2$, $10^3$, $10^4$, $10^5$, and $10^6$ CFU/insect (n=30 per treatment) and their survival was monitored over the course of seven days. PBS-injected larvae were used as control.

**Table S1. Generalized linear mixed model results for the effects of *TmRelish* knockdown and infection on AMP expression.** Results of generalized linear mixed models testing the effects of *TmRelish* knockdown, infection and their interaction on AMP expression, separately for each AMP.

| Gene name | Predictor | estimate | SE | Z value | *p* value |
|---|---|---|---|---|---|
| ***TmAtt1a*** | **kd. treatment** | -7.587 | 0.774 | -9.806 | **< 0.001** |
| | **infection** | -10.988 | 0.774 | -14.203 | **< 0.001** |
| | **kd. treatment: infection** | 7.802 | 1.094 | 7.131 | **< 0.001** |
| ***TmAtt1b*** | **kd. treatment** | -5.571 | 0.909 | -6.130 | **< 0.001** |
| | **infection** | -11.971 | 1.538 | -7.783 | **< 0.001** |
| | **kd. treatment: infection** | 4.885 | 2.175 | 2.246 | **0.025** |
| ***TmAtt2*** | **kd. treatment** | -3.808 | 0.7162 | -5.317 | **< 0.001** |
| | **infection** | -9.202 | 0.7162 | -12.848 | **< 0.001** |
| | **kd. treatment: infection** | 0.4937 | 1.0129 | 0.487 | 0.626 |
| ***TmTen1*** | **kd. treatment** | -3.7717 | 0.463 | -8.140 | **< 0.001** |
| | **infection** | -7.7504 | 0.710 | -10.916 | **< 0.001** |
| | **kd. treatment: infection** | 2.2323 | 1.004 | 2.223 | **0.026** |
| ***TmTen2*** | **kd. treatment** | -9.227 | 1.618 | -5.702 | **< 0.001** |
| | **infection** | -12.329 | 1.618 | -7.619 | **< 0.001** |
| | **kd. treatment: infection** | 8.011 | 2.288 | 3.501 | **< 0.001** |
| ***TmTen4*** | **kd. treatment** | -6.091 | 0.512 | -11.887 | **< 0.001** |
| | **infection** | -9.943 | 0.512 | -19.407 | **< 0.001** |
| | **kd. treatment: infection** | 5.588 | 0.725 | 7.712 | **< 0.001** |
| ***TmColA*** | **kd. treatment** | -5.973 | 0.941 | -6.346 | **< 0.001** |
| | **infection** | -12.560 | 0.941 | -13.344 | **< 0.001** |
| | **kd. treatment: infection** | 5.345 | 1.331 | 4.016 | **< 0.001** |
| ***TmColB*** | **kd. treatment** | -4.844 | 0.582 | -8.325 | **< 0.001** |
| | **infection** | -10.314 | 0.582 | -17.724 | **< 0.001** |
| | **kd. treatment: infection** | 4.131 | 0.822 | 5.020 | **< 0.001** |
| ***TmCec2*** | kd. treatment | -0.816 | 0.522 | -1.562 | 0.118 |
| | **infection** | -1.525 | 0.522 | -2.918 | **0.003** |
| | kd. treatment: infection | 0.996 | 0.739 | 1.349 | 0.177 |
| ***TmDef_L*** | **kd. treatment** | -3.580 | 0.647 | -5.535 | **< 0.001** |
| | **infection** | -7.879 | 0.647 | -12.181 | **< 0.001** |
| | kd. treatment: infection | 0.514 | 0.915 | 0.562 | 0.574 |

**Table S2. Generalized linear mixed model results for the effects of gut microbiota and infection on AMP expression**. Results of a generalized linear mixed models testing the effect of presence of gut microbiota, infection and their interaction on AMP expression, separately for each AMP.

| Gene name | Predictor | estimate | SE | Z value | *p* value |
|---|---|---|---|---|---|
| ***TmAtt1a*** | **gut microbiota** | 1.754 | 0.565 | 3.103 | **0.001** |
| | **infection** | -9.006 | 0.434 | -20.730 | **< 0.001** |
| | gut microbiota: infection | -1.166 | 0.614 | -1.898 | 0.058 |
| ***TmAtt1b*** | **gut microbiota** | 3.950 | 1.921 | 2.056 | **0.040** |
| | **infection** | -6.784 | 2.173 | -3.122 | **0.002** |
| | **gut microbiota: infection** | -5.319 | 2.238 | -2.377 | **0.018** |
| ***TmAtt2*** | gut microbiota | 0.513 | 0.424 | 1.210 | 0.226 |
| | **infection** | -9.627 | 0.424 | -22.681 | **< 0.001** |
| | gut microbiota: infection | -0.746 | 0.600 | -1.243 | 0.214 |
| ***TmTen1*** | gut microbiota | 0.668 | 0.603 | 1.107 | 0.268 |
| | **infection** | -7.890 | 0.603 | -13.083 | **< 0.001** |
| | gut microbiota: infection | -1.620 | 0.853 | -1.899 | 0.058 |
| ***TmTen2*** | gut microbiota | 0.832 | 1.479 | 0.563 | 0.574 |
| | **infection** | -6.440 | 1.512 | -4.260 | **< 0.001** |
| | gut microbiota: infection | -0.774 | 2.054 | -0.377 | 0.706 |
| ***TmTen4*** | **gut microbiota** | 1.441 | 0.556 | 2.592 | **0.010** |
| | **infection** | -9.506 | 0.556 | -17.099 | **< 0.001** |
| | **gut microbiota: infection** | -1.736 | 0.786 | -2.208 | **0.027** |
| ***TmColA*** | gut microbiota | 0.718 | 0.603 | 1.191 | 0.234 |
| | **infection** | -10.630 | 0.603 | -17.627 | **< 0.001** |
| | gut microbiota: infection | -1.1530 | 0.853 | -1.352 | 0.176 |
| ***TmColB*** | **gut microbiota** | 1.293 | 0.649 | 1.991 | **0.047** |
| | **infection** | -9.529 | 0.649 | -14.679 | **< 0.001** |
| | **gut microbiota: infection** | -1.979 | 0.918 | -2.156 | **0.031** |
| ***TmCec2*** | gut microbiota | -0.623 | 0.571 | -1.089 | 0.276 |
| | **infection** | -1.467 | 0.571 | -2.567 | **0.010** |
| | gut microbiota: infection | 0.433 | 0.808 | 0.536 | 0.592 |
| ***TmDef_L*** | gut microbiota | 1.000 | 0.705 | 1.418 | 0.156 |
| | **infection** | -7.076 | 0.913 | -7.750 | **< 0.001** |
| | **gut microbiota: infection** | -2.926 | 0.997 | -2.935 | **0.003** |

### Table S3. Primer sequences used in this study

| Gene Name | Primer sequence |
|---|---|
| *TmAtt1a*_qPCR_Fw<br>*TmAtt1a*_qPCR_Rv | 5′-GAAACGAAATGGAAGGTGGA-3′<br>5′-TGCTTCGGCAGACAATACAG-3′ |
| *TmAtt1b*_qPCR_Fw<br>*TmAtt1b*_qPCR_Rv | 5′-GAGCTGTGAATGCAGGACAA-3′<br>5′-CCCTCTGATGAAACCTCCAA-3′ |
| *TmAtt2*_qPCR_Fw<br>*TmAtt2*_qPCR_Rv | 5′-AACTGGGATATTCGCACGTC-3′<br>5′-CCCTCCGAAATGTCTGTTGT-3′ |
| *TmTen1*_qPCR_Fw<br>*TmTen1*_qPCR_Rv | 5′-CAGCTGAAGAAATCGAACAAGG-3′<br>5′-CAGACCCTCTTTCCGTTACAGT-3′ |
| *TmTen2*_qPCR_Fw<br>*TmTen2*_qPCR_Rv | 5′-CAGCAAAACGGAGGATGGTC-3′<br>5′-CGTTGAAATCGTGATCTTGTCC-3′ |
| *TmTen4*_qPCR_Fw<br>*TmTen4*_qPCR_Rv | 5′-GGACATTGAAGATCCAGGAAAG-3′<br>5′-CGGTGTTCCTTATGTAGAGCTG-3′ |
| *TmColA*_qPCR_Fw<br>*TmColA*_qPCR_Rv | 5′-GGACAGAATGGTGGATGGTC-3′<br>5′-CTCCAACATTCCAGGTAGGC-3′ |
| *TmColB*_qPCR_Fw<br>*TmColB*_qPCR_Rv | 5′-CAGCTGTTGCCCACAAGTG-3′<br>5′-CTCAACGTTGGTCCTGGTGT-3′ |
| *TmCec2*_qPCR_Fw<br>*TmCec2*_qPCR_Rv | 5′-TACTAGCAGCGCCAAAACCT-3′<br>5′-CTGGAACATTAGGCGGAGAA-3′ |
| *TmDefL*_qPCR_Fw<br>*TmDefL*_qPCR_Rv | 5′-GGGATGCCTCATGAAGATGTAG-3′<br>5′-CCAATGCAAACACATTCGTC-3′ |
| *TmL27a*_qPCR_Fw<br>*TmL27a*_qPCR_Rv | 5′-TCATCCTGAAGGCAAAGCTCCAGT-3′<br>5′-AGGTTGGTTAGGCAGGCACCTTTA-3′ |
| *TmRelish*_qPCR_Fw<br>*TmRelish*_qPCR_Rv | 5′-AGCGTCAAGTTGGAGCAGAT-3′<br>5′-GTCCGGACCTCATCAAGTGT-3′ |
| *dsTmRelish*_Fw<br>*dsTmRelish*_Rv | 5′-TAATACGACTCACTATAGGGGACGTGCACCATCAATA-3′<br>5′-TAATACGACTCACTATAGGGGCGTGTTTGGCCTTGAT-3′ |
| ds*EGFP*_Fw | 5′-TAATACGACTCACTATAGGGCTTAATGCACCACCACCACCAC-3′ |
| ds*EGFP*_Rv | 5′-TAATACGACTCACTATAGGGGTGACCCAGGATGTTACCGTC-3′ |
| 515F-16srRNA | 5′-GTGYCAGCMGCCGCGGTA-3′ |
| 806R-16srRNA | 5′-GGACTACNVGGGTWTCTAAT-3′ |

**※Underline indicates T7 promoter sequences.**
